# Autophagy Induced by Palmitic Acid: a Brake in NAFLD Neutrophils

**DOI:** 10.1101/2021.04.02.438261

**Authors:** Zhicheng Peng, Heyuan Wang, Alan Y. Hsu, Xiliang Du, Yuchen Yang, Baochen Fang, Yunfei Li, Yiwei Zhu, Yuxiang Song, Xiaobing Li, Zhe Wang, Xinwei Li, Guowen Liu

## Abstract

Innate immune suppression and high blood fatty acid levels are the pathological basis of multiple metabolic diseases. Neutrophil vacuolation is an indicator of the immune status of patients, which is associated with autophagy-dependent granule degradation. Vacuolated neutrophils are observed in ethanol toxicity and septicemia patients due to the changes in their blood constituents, but how about the neutrophils in nonalcoholic fatty liver disease (NAFLD) patient is unknown. Here, we confirmed that an adhesion deficiency and an increased autophagy level existed in NAFLD neutrophils, and the three neutrophil granule subunits, namely, the azurophil granules, specific granules and gelatinase granules, could be engulfed by autophagosomes for degradation, and these autophagy-triggered granule degradation events were associated with vacuolation in palmitic acid (PA)-treated and NAFLD neutrophils. Concordantly, the adhesion-associated molecules CD11a, CD11b, CD18 and Rap1 on the three granule subunits were degraded during PA induced autophagy. Moreover, the cytosolic CD11a, CD11b, CD18 and Rap1 were targeted by Hsc70 and then delivered to lysosomal-like granules for degradation. Notably, *in vitro and ex vivo*, PA induced autophagy by inhibiting the p-PKCα/PKD2 pathway. Overall, we showed that high blood PA level inhibited the p-PKCα/PKD2 pathway to induce NAFLD neutrophil autophagy, which promoted the degradation of CD11a, CD11b, CD18 and Rap1 and further decreased the adhesion of neutrophils, thereby impairing the neutrophil function of NAFLD patients. This theory provides a new therapeutic strategy to improve the immune deficiency in NAFLD patients.

**Visual Abstract:** 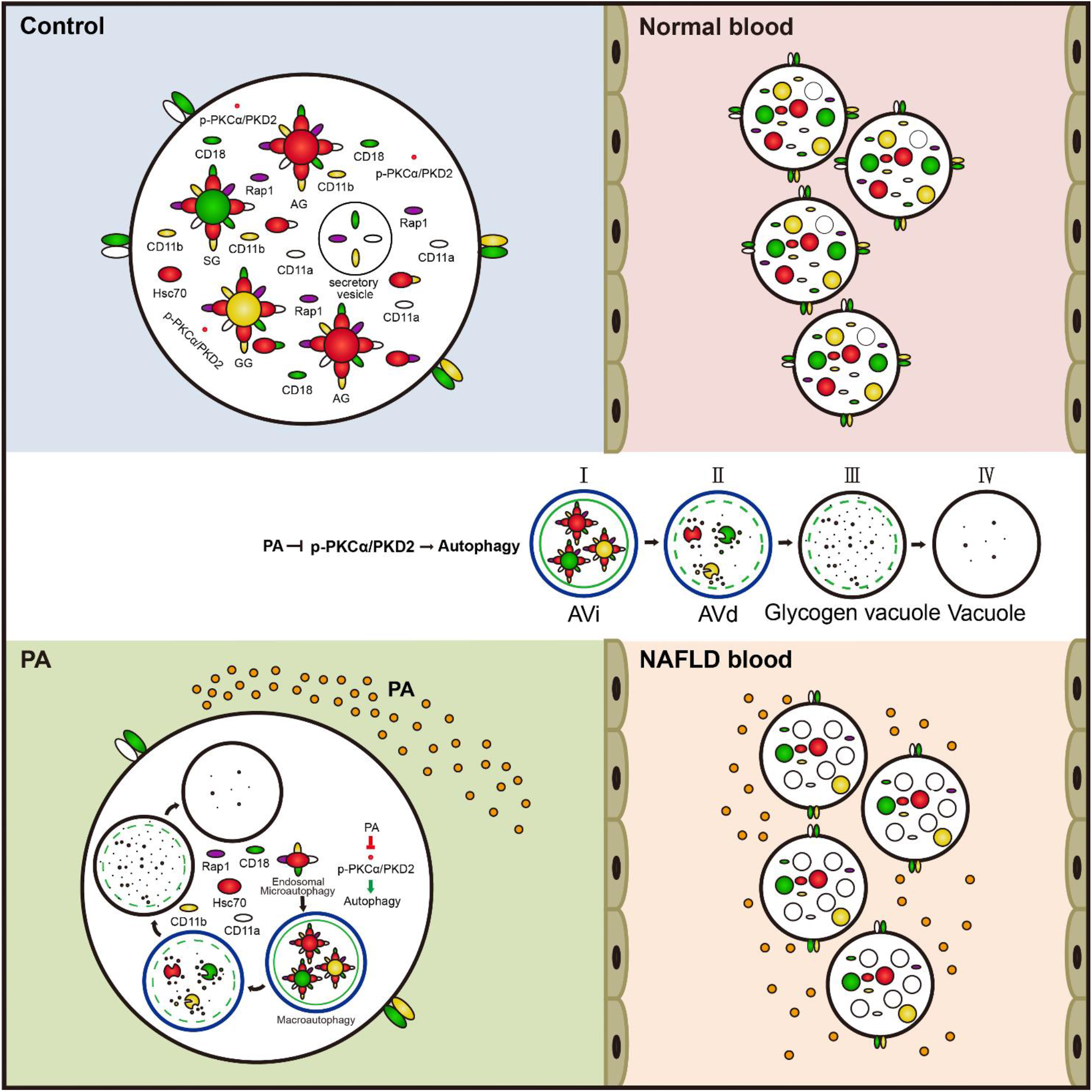

**Key Points:**

1. Vacuolation and adhesion deficiency of NAFLD neutrophils are associated with autophagy-dependent granule degradation
2. PA inhibits p-PKCα/PKD2 to induce autophagy, which induces the degradation of CD11a, CD11b, CD18 and Rap1 and decreases neutrophil adhesion

## Introduction

Neutrophils are the most abundant immune cells in human circulation, ranging from 40 to 70% of our circulating leukocytes^1^. Generally, neutrophils roll along the vessel walls to conduct immune surveillance. When they sense a chemotactic cue, the circulating neutrophils roll slowly to adhere to the venular endothelium, extravasate from the bloodstream and are rapidly recruited to infectious sites to provide the first line of defense against invading pathogens. Once there, neutrophils phagocytose the pathogen, release their anti-microbial content normally kept intracellularly in granules, and can release neutrophil extracellular traps to kill and prevent the dissemination of microbes. There are three neutrophil granules types: azurophil granules (AGs), specific granules (SGs) and gelatinase granules (GGs), each of which contains a specific array of microbicidal proteins, adhesion molecules, and various enzymes^2^.

In order for neutrophils to efficiently migrate to the site of infection, they express numerous adhesion molecules. Many β2 integrins^3^, including CD11a/CD18, CD11b/CD18, CD11c/CD18 and CD11d/CD18, are differentially expressed in neutrophils, either spatially or temporally (located in granules, plasma membranes and trafficked vesicles). These integrins mediate the firm adhesion between neutrophils and the endothelium by binding with intercellular adhesion molecule-1 (ICAM) and -2 after neutrophil activation^4–6^. CD11a/CD18 and CD11b/CD18 are the most abundant and critical β2 integrins during this process^7^. The small GTPase Rap1 is a key β2 integrin activity regulator^8^. Impaired Rap1 or excessive degradation of the β2 integrin subunits (α-chain or β-chain) leads to neutrophil adhesion and diapedesis deficiency, inducing a decrease in immune function characterized by persistent infections^9,10^.

Autophagy is a lysosome-dependent degradation process by which complete organelles (mitochondria) or other cytosolic cargoes are encapsulated and then delivered to lysosomes for degradation^11^. Previously, neutrophil vacuolation was associated with autophagy-triggered intracellular granule fusion events^12^. Interestingly, lysosome-associated membrane proteins (LAMPs), the autophagic receptor p62 and proteolytic enzymes were localized on AGs, SGs, and GGs^2^, suggesting that the three granule subunits of neutrophils are analogous to classic lysosomes^2,13^. These fusion events are speculated to be associated with p62-mediated autophagy-dependent granule degradation. Importantly, adhesion-associated molecules, such as β2 integrins and Rap1, were observed on the three granule subunits, and might be degraded together with granules.

Previous studies have demonstrated that ubiquitylated β1 integrins are sorted into early endosomes (EEs) or multivesicular bodies (MVBs) and then delivered to lysosomes for degradation via the endosomal sorting complex required for transport (ESCRT) machinery^5,14,15^. Furthermore, Rab GTPases (Rabs) and heat shock cognate 71 kDa protein (Hsc70) have been shown to mediate the fusion of EEs and MVBs with autophagosomes^16–18^, while integrins were located on Rab-positive EEs and MVBs^19,20^. In addition, molecular chaperone proteins, β2 integrins and Rap1 were found on the three lysosome-like granules^2^. Accordingly, we speculated that chaperone-mediated autophagy (CMA) mediated the degradation of ubiquitylated β2 integrins and Rap1 in neutrophils and further influenced neutrophil adhesion and migration.

Nonalcoholic fatty liver disease (NAFLD) is a highly prevalent condition which affects 25% of the population worldwide. NAFLD has been associated with obesity, insulin resistance, type 2 diabetes mellitus (T2DM), hypertension, hyperlipidemia, and metabolic syndromes. High blood palmitic acid (PA) levels are a major pathological hallmark of NAFLD and have been shown to be a direct activator of autophagy via the downregulation of protein kinase C α subunit (PKCα)^21^. A recent study revealed that knockout or pharmacological inhibition of PKCα dramatically increased autophagy^22,23^, suggesting that PKCα is a negative regulator of autophagy. We speculated that in NAFLD patients, when cells are exposed to high blood concentrations of PA, autophagy in neutrophils could contribute to dampening neutrophil function, thus alleviating inflammatory functions^21^. However, the mechanism via which PA influences neutrophil autophagy, adhesion and diapedesis in NAFLD patients is unknown.

Here, we identified a mechanism in which the three neutrophil granule subsets of neutrophils were degraded by autophagy. *Ex vivo* and in vitro, PA treatment induced autophagy via the p-PKCα/PKD2 pathway, decreasing the accumulation of CD11a, CD11b, CD18 and Rap1, and leading to deficiencies in neutrophil adhesion and diapedesis. In summary, here we present evidence that autophagy plays a bridging role between metabolic diseases, such as fatty liver disease and deficiency in neutrophil adhesion and diapedesis.

## Materials and methods

### Antibodies and reagents

Anti-CD11a mAb (Ab52895), Anti-CD11b mAb (Ab52478), Anti-CD11b mAb (Ab34216), Anti-CD18 mAb (Ab53009), Anti-CD18 mAb (Ab657), Anti-Hsc70 mAb (Ab2788), Anti-Hsc70 mAb (ab51052), Anti-Rap1 mAb (Ab175329), Anti-LC3A/B Ab (Ab128025), Anti-SQSTM1/p62 Ab (Ab101266), Anti-SQSTM1/p62 Ab (Ab56416), Anti-Myeloperoxidase mAb (Ab25989), Anti-Lactoferrin Ab (Ab112968), Anti-PKCα (phospho T638) mAb (Ab32502) were purchased from Abcam (Cambridge, MA, USA). Anti-ATG5 Ab (NB110-53818), BafilomycinA1 (CAS88899-55-2) were purchased from Novus Biologicals (Centennial, CO, USA). Anti-Ubiquitin (K-48) mAb (sc-271289), Anti-PKD2 mAb (sc-374344), Anti-β-actin mAb (sc-47778) were purchased from Santa Cruz Biotechnology (Dallas, TX, USA). Anti-rabbit IgG conjugated to 5-nm Gold (G7277), anti-mouse IgG conjugated to 10-nm Gold (G7777), anti-goat IgG conjugated to 10-nm Gold (G5402), anti-mouse IgG conjugated to 5-nm Gold (G7527) were purchased from Sigma Aldrich (Shanghai, China). Anti-MMP-9 mAb (MA5-15886), Anti-CD11a mAb (MA1-19003) were purchased from Thermo Scientific (Waltham, MA, USA). Sodium palmitate (P9767), Chloroquine phosphate (PHR1258), MG132 (M8699), fMLP (F3506), DFP (D0879) were purchased from Sigma Aldrich (Shanghai, China). Human myeloperoxidase ELISA Kit was purchased from Jianglai Biotechnology (Shanghai, China). Cell Fractionation Kit (9038) was purchased from Cell Signaling Technology (Danvers, MA, USA).

### Preparation of the PA/BSA Complex Solution

Sodium palmitate was dissolved in distilled water by heating at 70°C till completely dissolve. Simultaneously, 10% (wt/vol) FFA-free BSA solution was prepared at 55°C. The two solutions were mixed and coupled at 55°C for 10 min, made into 50 mM of PA/BSA complex stock solution. Equal volume 5% (wt/vol) FFA-free BSA solution treatment as control.

### Human

Venous blood and liver samples (normal and NAFLD) were collected from the First Hospital of Jilin University. Written informed consent was obtained from all subjects in compliance with the Declaration of Helsinki guidelines and approved by the ethics committee of the First Hospital of Jilin University (2016-416). Subjects with other causes of chronic liver disease and renal dysfunction or those receiving potentially hepatotoxic drugs were excluded.

### HL-60

The HL-60 cell line was purchased from Keygentec (Nanjing, China) and was cultured in IMDM medium supplemented with 20% FBS. The HL-60 cells were differentiated to a neutrophil-like phenotype with a final concentration of 1.3% DMSO for 6 d.

### Preparation of Neutrophils

Normal and NAFLD neutrophils were isolated using commercialized kit purchased from Tbdscience (Tianjin, China) according to the manufacturer’s protocol. Neutrophils were cultured in RPMI 1640 complete medium. Neutrophil viability was higher than 97% as assessed by trypan blue staining, and purity was higher than 98% as analyzed by Wright and Giemsa staining.

### Neutrophil Viability Assays

Neutrophil were seeded in the 96-wells cell culture plate at a density of 1.0×10^6^ cells/mL and cultured with RPMI 1640 containing 5% (wt/vol) FFA-free BSA with or without PA (0.25 mM) for various time points. Neutrophil viability was determined by Cell Counting Kit-8 assay (CK04, Tongren, Japan) according to the manufacturer’s protocol. OD value was measured using a microplate reader at wavelength in the 570 nm, which reflected the viability of neutrophils.

### Transmission Electron Microscopy

Cells were pelleted and fixed with 4% glutaraldehyde in 0.1 M PBS overnight at 4°C. Subsequently, postfixed in 1% osmium tetroxide was followed by dehydration with graded series of ethanol, infiltration and embedding in SPI-PON 812 resin (SPI Supplies, West Chester, PA, USA). Ultrathin sections with a thickness of 65 nm were cut using a microtome Leica EM UC7 (Leica Microsystems Company, Wetzlar, Hessen, Germany) and poststained with 2% uranyl acetate for 10 min and 0.3% lead citrate for 10 min. The ultrathin sections were observed using a Hitachi H-7650 transmission electron microscope (Hitachi, Kyoto, Japan).

### Immunogold Electron Microscopy

Neutrophils were prepared for immunogold electron microscopy as previously described with minor modifications^24^. Briefly, neutrophils were fixed in 4% paraformaldehyde and 0.5% glutaraldehyde at 4°C for 1.5 h, washed, scraped and pelleted, sectioned. The sections were soaked in pure water and then blocked in 3% skimmed milk in PBS for 30 min at room temperature. Subsequently, the sections were labeled with primary antibodies followed by secondary antibodies conjugated with protein A-gold. The sections were poststained with 2% uranyl acetate for 10 min before observation with a Hitachi H-7650 transmission electron microscope (Hitachi, Kyoto, Japan).

### Granule quantification

The granules number were performed using the transmission electron microscopy. The sections were chosen by randomly and then the granule number of each section was quantified based on their morphological characteristics, such as the size, shape and electron density: AGs are the largest granules (the average diameters approximately 200 nm), with spherical and ellipsoid 2 kind of shape and high electron density; SGs are smaller than AGs, dumbbell shape and with lower electron density; GGs are the smallest granule in size, round shape and with the lowest electron density^25,26^. However, duo to the thickness of the sections is approximately 90 nm, which is thinner than the diameter of AGs. So, the density of the AGs we observed much lighter than SGs^26^. In addition, SGs sometimes show the round shape duo to the different cut angle or cut ways.

### Marker Assays

Granules can be distinguished on the basis of their morphological characteristics, such as size, shape and electron density: AGs are large granules with high electron density; SGs are smaller than AGs and have lower electron densities; and GGs are the smallest granules in size and have the lowest electron densities^27^. In addition, the three granule subunits are identifiable by their marker proteins: myeloperoxidase (MPO) is an AG marker, lactoferrin is an SG marker, and gelatinase (MMP-9) is a GG marker^28–30^. These marker molecules were labeled using immunogold electron microscopy.

### Adhesion Assay

Human umbilical vein endothelial cells (HUVECs, KG060) were purchased from Keygentec (Nanjing, China) and were cultured in RPMI 1640 medium supplemented with 10% FBS. The monolayer of HUVECs were plated 12h ahead in 96-well plates as substrates. Then, the neutrophils or HL-60 cells were collected after different treatments. Firstly, the cells were labeled with calcein AM (5 μM) for 30 min at 37°C. Subsequently, the labeled cells were washed, resuspended at 5.0×10^6^ cells/mL, and then were activated with 1 µM fMLP for 30 min, and then added to the HUVEC monolayer. The plates were incubated at 37°C for 30-60 min. Notably, for ex vivo experiments, neutrophils from NAFLD patients or normal subjects were incubated with their own sera at this step. At the end of the incubation period, nonadherent cells were removed with cold RPMI medium containing 1% FBS. The plate was scanned with a Tecan Infinite 200 PRO multifunctional microplate reader (TECAN, Männedorf, Switzerland), and the fluorescence of the adherent cells was measured by Nikon fluorescence microscope (Nikon, Tokyo, Japan).

### Immunoprecipitation Assay

Neutrophils were harvested and incubated with cold 1×PBS containing 2 mM DFP on ice for 15 min. Cytosolic fraction of neutrophils was performed using a Cell Fractionation Kit (Danvers, MA, USA). Immunoprecipitated was performed from cytosolic fraction using a Pierce Crosslink Immunoprecipitation Kit (Thermo Scientific, MA, USA). 1 mg of cytosolic fraction was precleared with 80 µL of the control agarose resin slurry for 1.5 h at 4°C. The primary antibodies were cross-linked to protein A/G plus agarose. The precleared lysate was added to the primary antibody-crosslinked resin in the column overnight at 4°C. The unbound proteins were washed away with IP lysis/wash buffer. Then, the immunoprecipitated proteins were eluted. The eluate concentrations were determined using the BCA Protein Assay Kit (Pierce, IL, USA). The protein complexes were analyzed by SDS-PAGE, and the gel was stained with Coomassie blue.

### Generation and differentiation of the ATG5, p62 and Hsc70 knockdown and PRKD2 overexpression HL-60 cell lines

The lentiviral vectors for ATG5 knockdown (LV-GFP-shATG5), p62 knockdown (LV-GFP-shp62) and for Hsc70 knockdown (LV-GFP-shHsc70) were purchased from GeneChem (Genechem, Shanghai, China). The lentiviral vector for PRKD2 overexpression (LV-GFP-PRKD2) was constructed by GeneChem (Genechem, Shanghai, China). HL-60 cells were infected with lentiviral vectors at a MOI of 25 in the presence of 5 μg/mL polybrene. The HL-60 cells were differentiated to a neutrophil-like phenotype with a final concentration of 1.3% DMSO for 6 d. Stably knocking down or overexpressing cell lines were selected with puromycin (5 μg/ml) and identified by qRT-PCR, western blotting and immunofluorescence. The wild-type and the relevant empty lentivectors cells were used as negative control.

### Protein Complex Identification by Shotgun Analysis

Endogenous CD11a, CD11b, CD18, Rap1, and Hsc70 were enriched using immunoprecipitation assay. Approximately 30 μg of IP complexes of CD11a, CD11b, CD18, Rap1, and Hsc70 was performed by Shotgun Analysis as previously described^31^. MS/MS spectra were searched using MASCOT engine (version 2.2, Matrix Science) embedded into Proteome Discoverer 1.4^32^ against the Uniprot Human database (156914 sequences, downloaded on March 2, 2017). For protein identification, the following parameters were selected: Peptide mass tolerance: 20 ppm, MS/MS tolerance: 6 ppm, Enzyme: Trypsin, Max Missed Cleavages: 2, Fixed modifications: Carbamidomethyl (C), Dynamical modifications: Oxidation (M) and GlyGly (K), peptides FDR ≤ 0.01, protein FDR ≤ 0.01, Filter by score ≥ 20.

### Isobaric Tag for Relative and Absolute Quantitation (iTRAQ) Proteomic Assay

Neutrophils were treated with PA (0.25 mM) for 3 h, and the untreated group served as control. The samples were lysed with SDT buffer (4%SDS, 100mM Tris-HCl, 1mM DTT, pH7.6) completely and centrifuged at 14,000×g at room temperature for 5 min. Then, the supernatants were collected, and the protein concentrations were determined using the BCA Protein Assay Kit (Pierce, IL, USA). Twenty micrograms of protein were separated on a 12.5% SDS-PAGE gel (with a constant current of 14 mA for 90 min) to evaluate protein quality. Protein bands were visualized with Coomassie blue R-250 staining. The process of Trypsin Digestion, iTRAQ Labeling, Peptide Fractionation and LC-MS/MS analysis were performed as previously described^33^. MS/MS spectra were searched using MASCOT engine (version 2.2, Matrix Science) embedded into Proteome Discoverer 1.4^32^ against the Uniprot Human database (156639 sequences, downloaded on January 5, 2017). For protein identification and quantification, the parameters were selected as previously described^33^. The median protein ratio should be 1 after the normalization.

### Quantification and statistical analysis

Statistical analysis was carried out using PRISM 6 (GraphPad). The unpaired t-test (when comparing 2 groups), One-way ANOVA test (when comparing with a single group), and two-way ANOVA (when comparing multiple factors between two groups) were used in this study as indicated in the figure legends. Individual P values are indicated in the figures, with no data points excluded from statistical analysis.

## Results

### Autophagy-dependent Vacuolation and Adhesion Deficiency Existed in NAFLD Neutrophils

Neutrophil vacuolation is an indicator of the immune status of patients^34^. An increase in vacuolated neutrophils are commonly observed in ethanol toxicity and septicemia patients^35,36^, which have been associated with autophagy-triggered granule degradation^12,37^, Whether vacuolization of neutrophils also contribute in NAFLD patients is currently unknown. To investigate the immune status of NAFLD patients, neutrophils were obtained from normal individuals (n=12) and NAFLD patients (n=12). NAFLD patients were diagnosed by liver biopsy and hepatic HE staining (Figure S1). The clinical parameters of the subjects are listed in Table 1. Four consecutive stages of autophagic vacuoles in NAFLD neutrophils were defined, based on the degree of degradation of cytosolic portions or granules: early autophagic vacuoles (AVi), degradative autophagic vacuoles (AVd), glycogen vacuoles, and vacuoles. The four stages of autophagic vacuoles were observed in NAFLD neutrophils, with increased total vacuole number and autophagic vacuole to neutrophil area ratio, compared to normal neutrophils (Figure 1A and Figure 1B). The lipidation level of LC3B was also markedly increased and the accumulation of p62 decreased in NAFLD neutrophils (Figure 1C). These results indicated that the autophagic process was more prominent in NAFLD neutrophils. Interestingly, the number of granules in NAFLD neutrophils was significantly reduced (Figure 1D). In the AVi and AVd stages, a significant number of granules were engulfed by autophagic vacuoles (Figure 1A). To evaluate whether granule-associated adhesion molecules were degraded with granules in NAFLD neutrophils, the total protein levels and the surface expression of CD11a, CD11b CD18 and Rap1 were assessed by immunoblotting and by flow cytometry, respectively. As expected, the total protein level and surface expression of CD11a, CD11b and CD18 were all significantly lower in patients compared to the healthy controls (Figure 1C, Figure 1E and Figure S2). The reduced expression of the adhesion molecules correlated with an impaired level of adhesion of NAFLD neutrophils in vitro compared to healthy neutrophils (Figure 1F). These results showed that NAFLD neutrophils had an increased number of autophagic vacuoles correlated to a decreased adhesion deficiency, suggesting that autophagy inhibited the motility of neutrophils in NAFLD patients.

**Table 1.** The anthropometric, biochemical parameters and clinical characteristics of normal individuals and NAFLD patients

| Parameter | Normal (n=12) | NAFLD (n=12) | P-value |
| --- | --- | --- | --- |
| Gender (female) | 6 (50.0%) | 6 (50.0%) |  |
| Age (years) | 36±15 | 46±10 | 0.007 |
| Body weight (kg) | 67.1 | 77.7 | 0.020 |
| Height (m) | 172 | 169 | 0.210 |
| Non-esterified fatty acids (mmol/L) | 0.35±0.042 | 0.53±0.028 | 0.002 |
| Fasting palmitic acid (mmol/L) | 0.094±0.008 | 0.19±0.007 | 0.000 |
| BMI (kg/m2) | 22.54±0.34 | 27.02±0.86 | 0.000 |
| Fasting glucose (mmol/L) | 5.07±0.20 | 5.37±0.18 | 0.271 |
| TC (mmol/L) | 4.82±0.18 | 5.14±0.36 | 0.437 |
| TG (mmol/L) | 1.37±0.11 | 1.89±0.29 | 0.110 |
| HDL cholesterol (mmol/L) | 1.39±0.13 | 1.11±0.08 | 0.072 |
| LDL cholesterol (mmol/L) | 2.91±0.27 | 3.14±0.25 | 0.550 |
| AST (U/L) | 21.83±1.13 | 30.75±5.55 | 0.130 |
| ALT (U/L) | 25.25±3.36 | 33.17±10.48 | 0.479 |
| AST/ALT | 0.99±0.11 | 1.15±0.12 | 0.323 |
| White blood cells (10<sup>9</sup>/L) | 6.02±0.54 | 6.86±0.50 | 0.267 |
| Neutrophils (10<sup>9</sup>/L) | 3.53±0.32 | 4.18±0.43 | 0.234 |
| Neutrophils/White blood cells (%) | 0.58±0.02 | 0.60±0.02 | 0.395 |
| Lymphocytes (10<sup>9</sup>/L) | 2.09±0.19 | 2.11±0.12 | 0.930 |
| Monocytes (10<sup>9</sup>/L) | 0.31±0.06 | 0.41±0.06 | 0.194 |

BMI, body mass index; TC, total cholesterol; TG, Triglycerides; HDL, high-density lipoprotein cholesterol; LDL, low density lipoprotein cholesterol; AST, aspartate aminotransferase; ALT, alanine aminotransferase.

**Figure 1.**
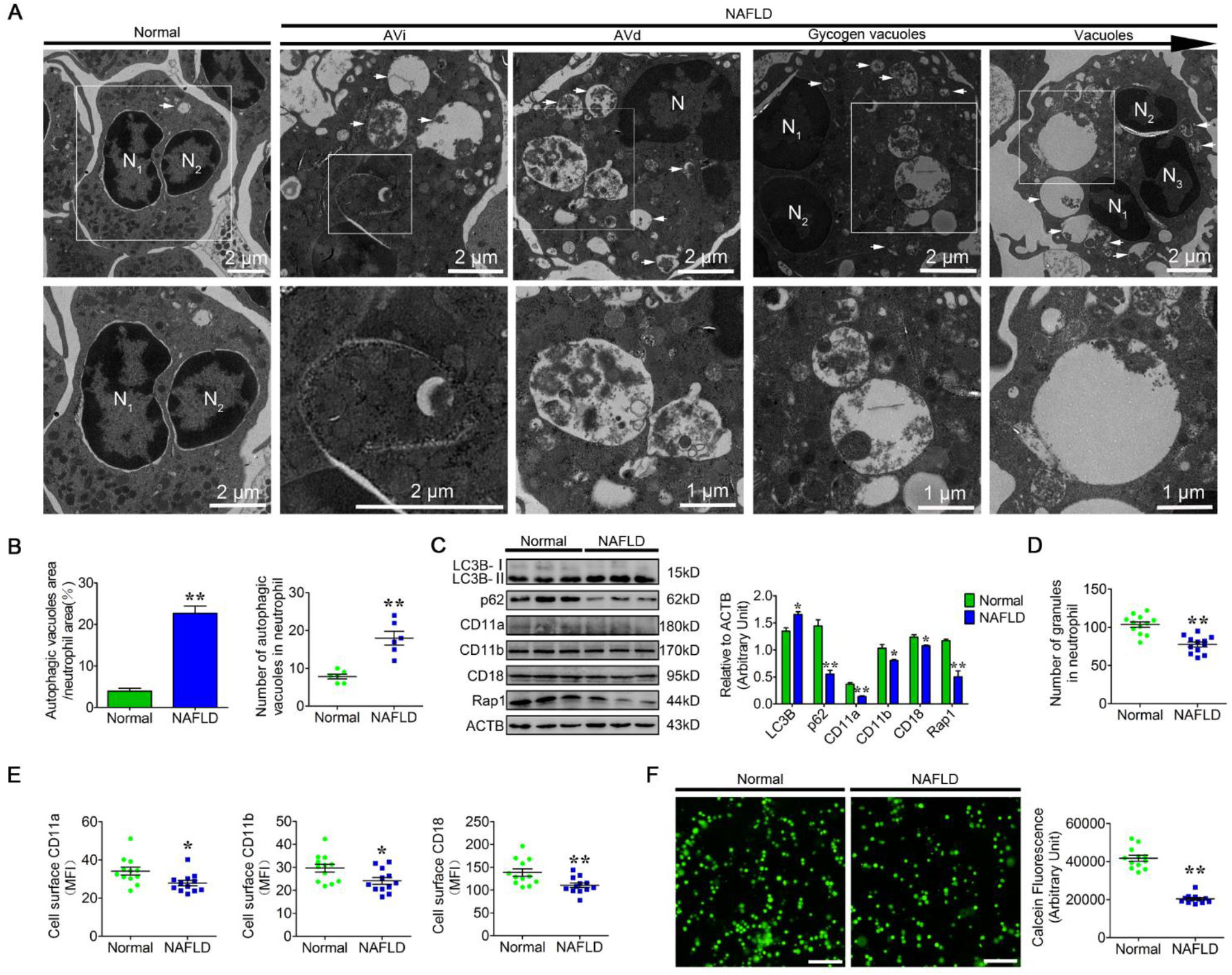
Autophagy-dependent Vacuolation and Adhesion Deficiency Existed in NAFLD Neutrophils. NAFLD neutrophils for transmission electron microscope were directly fixed after isolated from the patient’s blood without incubation, for adhesion assay, normal and NAFLD neutrophils were incubated with the subjects serum. (A) Representative transmission electron micrographs of normal and NAFLD neutrophils. White arrows indicate autophagic vacuoles (AVi, AVd, glycogen vacuoles and vacuoles). N (N1, N2, N3), nucleus. Scale bars as indicated. (B) The area ratio of autophagic vacuoles to neutrophils and the number of autophagic vacuoles in neutrophils were determined (n = 6). Data represent the mean ± s.e.m. (** P < 0.01 versus the control group; Significance calculated using t test). (C) Immunoblot for LC3B, p62, CD11a, CD11b, CD18, and Rap1 in normal and NAFLD neutrophils. ACTB was used as a loading control (n = 3). Data represent the mean ± s.e.m. (* P < 0.05 and ** P < 0.01 versus the control group; Significance calculated using two-way ANOVA). (D) The number of granules in normal and NAFLD neutrophils was determined (n = 12). Data represent the mean ± s.e.m. (** P < 0.01 versus the control group; Significance calculated using t test). (E) Surface expression of CD11a, CD11b and CD18 on normal and NAFLD neutrophils (n = 12). Surface expression of CD11a, CD11b and CD18 was assessed by flow cytometry analysis (n = 12). MFI, mean fluorescence intensity. Data represent the mean ± s.e.m. (* P < 0.05 and ** P < 0.01 versus the control group; Significance calculated using t test). (F) Representative fluorescence micrograph images (left) and fluorescence microplate analysis (right) of normal and NAFLD neutrophils adhered to HUVECs (n = 12). Scale bar, 400 μm. Data represent the mean ± s.e.m. (** P < 0.01 versus the control group; Significance calculated using t test).

### PA Enhanced Autophagy and Degraded the Granules in Neutrophils

PA, a major pathological hallmark of NAFLD, can induce autophagy in mouse embryonic fibroblasts^21^. We investigated the effect of 0.25 mM PA, a pathological concentration in NAFLD, on autophagy in neutrophils. Immunoblotting results showed that neutrophil autophagy was induced between 4 and 8 hours of PA treatment (Figure S3). PA also strongly triggered neutrophil vacuolation (Figure?). Consistent with our *ex vivo* findings, the four stages of autophagic vacuoles were observed in PA-treated neutrophils (Figure 2A). The number of autophagic vacuoles and the ratio of autophagic vacuole area to neutrophil area were significantly higher in PA-treated neutrophils than in control neutrophils (Figure 2B). The lipidation levels of LC3B were significantly higher in PA-treated neutrophils, while p62 were significantly reduced (Figure 2C). Furthermore, the number of granules significantly decreased in PA-treated neutrophils (Figure 2D). To make sure autophagy mediates the decrease of granules triggered by PA, Rapamycin (RAP) was used to activate autophagy. Expectedly, the number of granules decreased in RAP treated neutrophils (Figure 2D). Moreover, when autophagy induced by PA was blocked by bafilomycin A (BafA1) or hydroxychloroquine sulfate (CQ), the number of granules significantly increased (Figure 2D). However, PA induced neutrophil degranulation would also affect granule numbers (Figure S4A), To exclude this, neutrophils were stimulated with fMLP to further induce degranulation upon PA treatment. No difference was observed in granule numbers compared to the unstimulated groups (Figure S4B). These results suggests that PA induced autophagy plays an important role in neutrophil granule homeostasis. ATG5 knockdown lentivirus were infected the neutrophil-like differentiated (dHL-60) cells to deficient autophagy to further reiterate the effect of PA-induced autophagy. PA-treated ATG5-KD dHL-60 cells showed a similar recovery of granule number (Figure 2E). These results indicate that PA strongly enhances neutrophil autophagy and vacuolation, which is associated with granule degradation.

**Figure 2.**
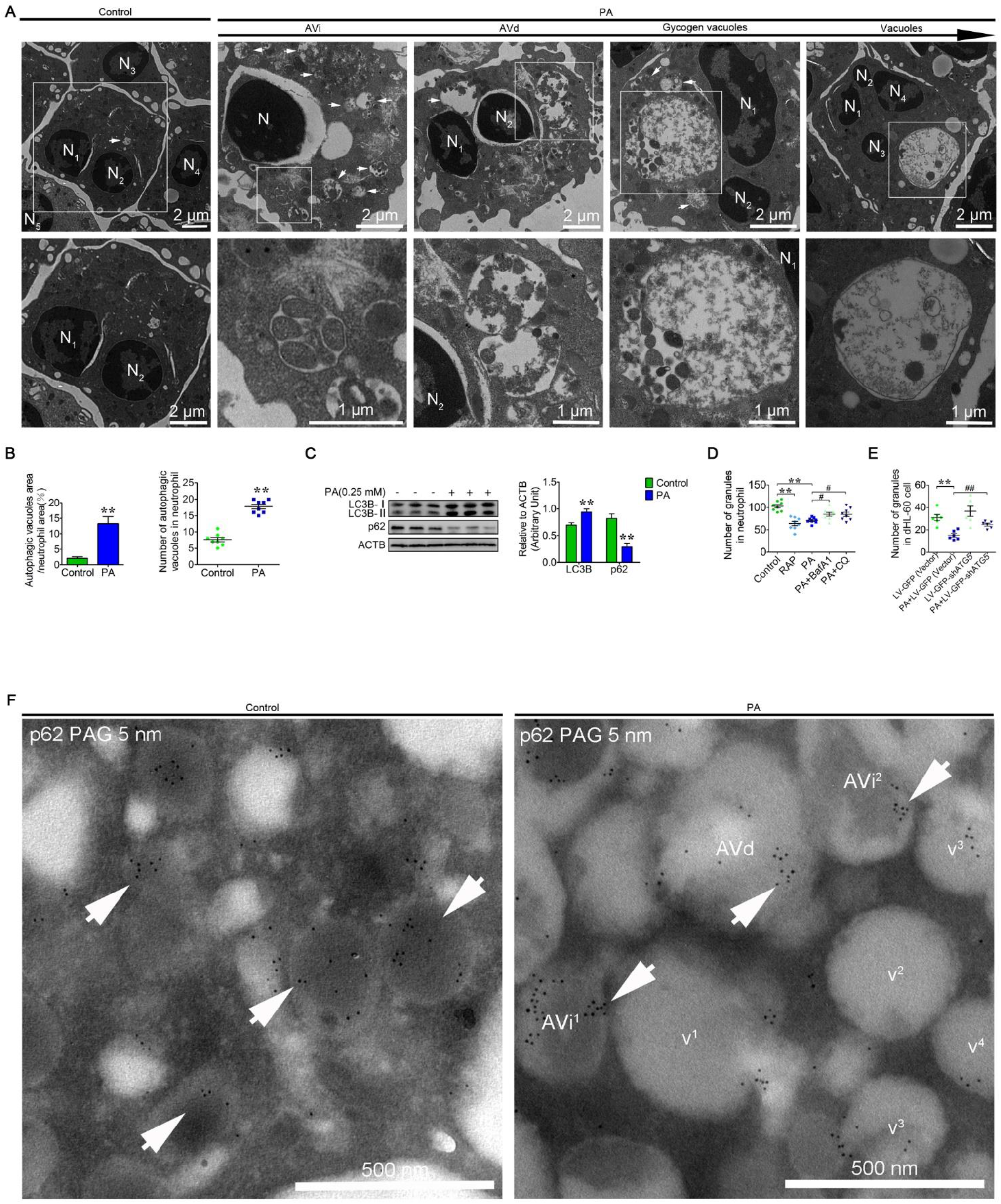
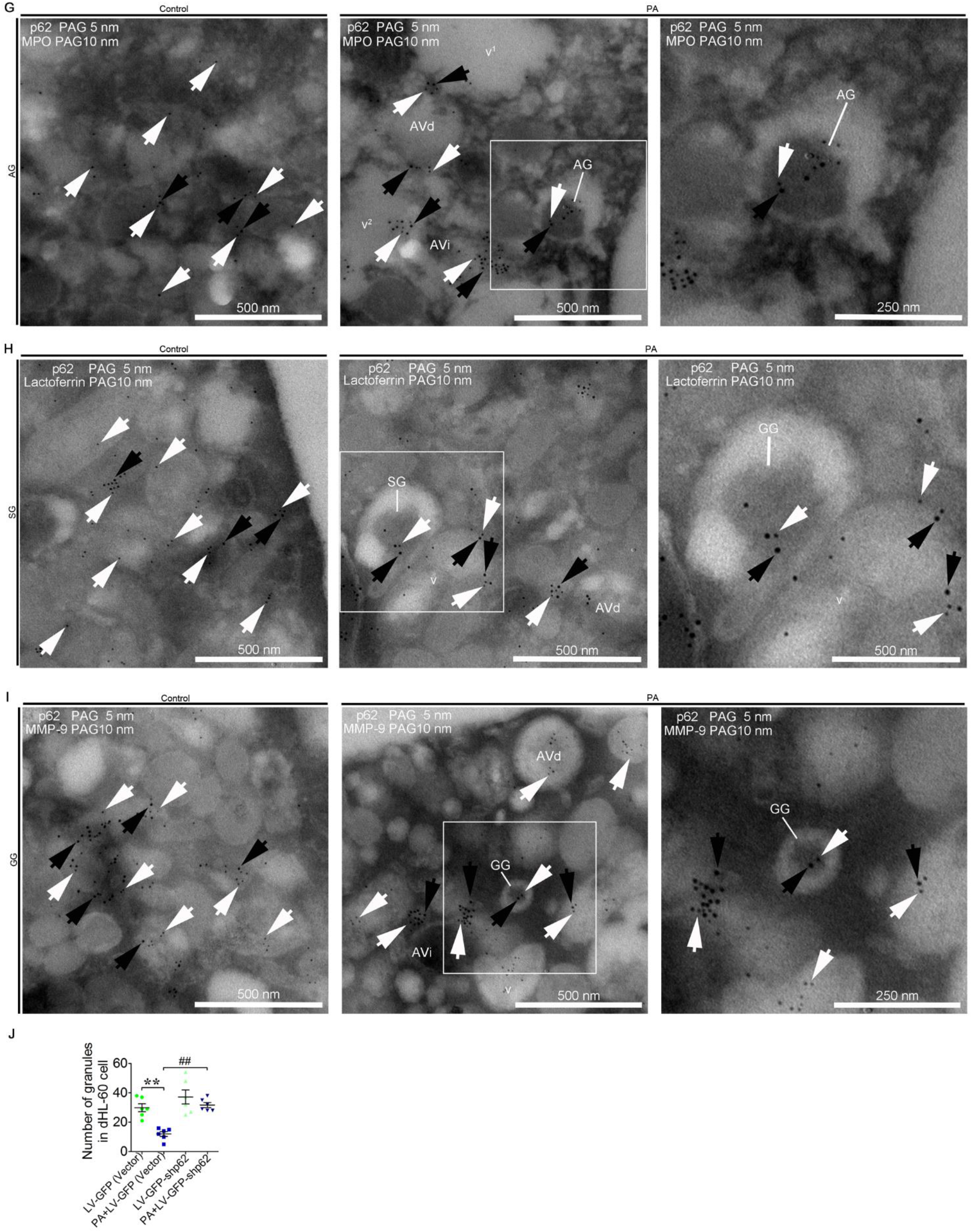
PA Enhanced Autophagy and Degraded Granules in Neutrophils. (A) Representative transmission electron micrographs of control and PA (0.25 mM)-treated neutrophils (left). White arrows indicate autophagic vacuoles (AVi, AVd, glycogen vacuoles and vacuoles). N (N1, N2, N3…), nucleus. Scale bars as indicated. (B) The area ratio of autophagic vacuoles to neutrophils and the number of autophagic vacuoles in neutrophils were determined (right, n = 8). Data represent the mean ± s.e.m. (** P < 0.01 versus the control group; Significance calculated using t test). (C) Immunoblot for LC3B and p62 in control and PA-treated neutrophils (n = 3). Data represent the mean ± s.e.m. (** P < 0.01 versus the control group; Significance calculated using two-way ANOVA). (D) The number of granules in control, RAP and PA-treated (treated or not treated with BafA1, CQ) neutrophils was determined (n = 8). Data represent the mean ± s.e.m. (** P < 0.01 versus the control group, # < 0.05 and versus the PA-treated group; Significance calculated using one-way ANOVA). (E) The number of granules in control and PA-treated dHL-60 cells (infected with LV-GFP-shATG5 or empty lentivectors) was determined (n = 6). Data represent the mean ± s.e.m (** P < 0.01 versus the control group, ## p < 0.01 versus the PA-treated group; Significance calculated using one-way ANOVA). (F) Immunogold electron micrograph showing the localization of p62 in control and PA-treated neutrophils. Five-nanometer p62 grains (white arrows) were observed on the granules (control) and autophagic vacuoles (PA). Scale bars as indicated. (G-I) Partial view of immunogold electron micrograph showing the colocalization of p62 with the AG marker MPO (G), SG marker lactoferrin (H) and GG marker MMP-9 (I) in control and PA-treated neutrophils. White arrows (5-nm gold grains) indicate p62. Black arrows (10-nm gold grains) indicate MPO, lactoferrin and MMP-9. v (v1, v2, v3…), vacuoles. Scale bars as indicated. (J) The number of granules in control and PA-treated dHL-60 cells (infected with LV-GFP-shp62 or empty lentivectors) was determined (n = 6). Data represent the mean ± s.e.m. (** P < 0.01 versus the control group, ## p < 0.01 versus the PA-treated group; Significance calculated using one-way ANOVA).

Organelles delivery to lysosomes for degradation is dependent on the autophagic receptor p62^38^. To investigate whether the decrease in granule levels was associated with p62-mediated granule degradation, immunogold electron microscopy was performed. The p62 electron-dense gold particles were predominantly localized on the AVi and AVd, but rarely on the vacuoles in PA-treated neutrophils (Figure 2F). This data motivated us to speculate that p62 might mediate the degradation of granules. AGs, SGs and GGs can be distinguished according to their size and electron density^27^. In addition, myeloperoxidase (MPO), lactoferrin and gelatinase (MMP-9) are markers of AG, SG, and GG, respectively^28–30^. To investigate whether all three granule types could be degraded by autophagy, double-labeling studies and morphological analysis were performed in control and PA-treated neutrophils. While p62 and MPO were colocalized on the large and highly electron-dense AGs (Figure 2G); p62 and lactoferrin were colocalized on the smaller and less electron-dense SGs (Figure 2H); and p62 and MMP-9 were colocalized on the smallest and least electron-dense GGs (Figure 2I). The AGs, SGs and GGs were observed could be engulfed by autophagic vacuoles (Figure 2G, Figure 2H and Figure 2I) Expectedly, knockdown of p62 attenuated the PA-induced decreased granule number in dHL-60 cells (Figure 2J). Altogether, this showed that p62 mediated the degradation of the AGs, SGs and GGs by autophagy, thereby causing the decrease in granule number and neutrophil vacuolation.

### PA-induced Autophagy Decreased Neutrophil Adhesion

In control neutrophils, CD11a, CD11b, CD18 and Rap1 electron-dense gold particles were present on the granules, secretory vesicles, and the plasma membrane as well as in the cytoplasmic matrix (Figure 3A). In PA-treated neutrophils, the CD11a, CD11b, CD18 and Rap1 gold particles were primarily present on AVi and AVd, but sparsely on vacuoles (Figure 3A). AGs, SGs and GGs were distributed with Rap1 in neutrophils^2^. We next investigated whether CD11a, CD11b and CD18 were also detected on AGs, SGs and GGs using double-labeling (CD11a, CD11b and CD18 colocalized with MPO, lactoferrin and MMP-9, respectively) and the morphological analysis of the three granule subtypes. We found that the gold particles corresponding to CD11a, CD11b and CD18 were all present on AGs (Figure 3B), SGs (Figure 3C) and GGs (Figure 3D), and few located on vacuoles (Figure 3B, Figure 3C and Figure 3D). These results suggested that the degradation of the three granules could possibly be mediated by autophagy. Consistent with this observation, the protein levels of CD11a, CD11b, CD18 and Rap1 were significantly lower in PA-treated neutrophils than in control neutrophils (Figure 3E). Furthermore, surface expression of CD11a, CD11b, and CD18 were greatly decreased in PA-treated neutrophils (Figure 3F and Figure S5). As decreased surface levels of CD11a, CD11b, and CD18 influence neutrophil adhesion, we measured neutrophil’s adhesion with or without PA n. Cells were seeded on collagen-coated culture plates. Control neutrophils adhered to the plates evenly and tightly (Figure S6), while PA-treated PMN adhered to the plates loosely, and cell lumps were observed floating in the medium (Figure S6). Quantification of the cell attachment showed that the adhesion of PA-treated neutrophils was significantly impaired compared to control neutrophils (Figure 3G). This was not due to a cytotoxic effect of PA on neutrophils at the concentration used in our experiments (0.25 mM) (Figure S7). Taken together, the results showed that PA induced autophagy triggered AG, SG and GG degradation and was accompanied by the degradation of CD11a, CD11b, CD18, and Rap1 in neutrophils, significantly decreasing neutrophil adhesion.

**Figure 3.**
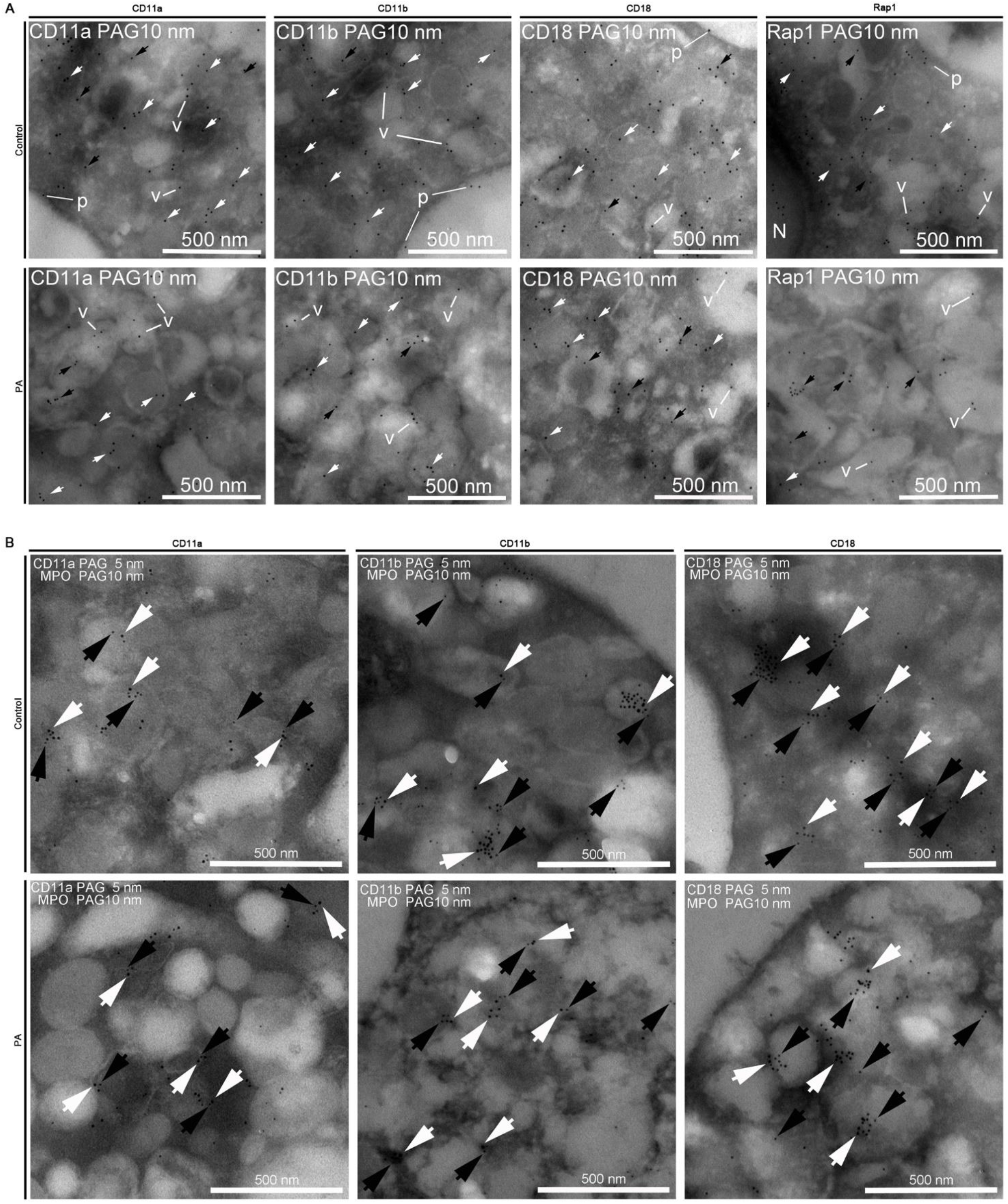
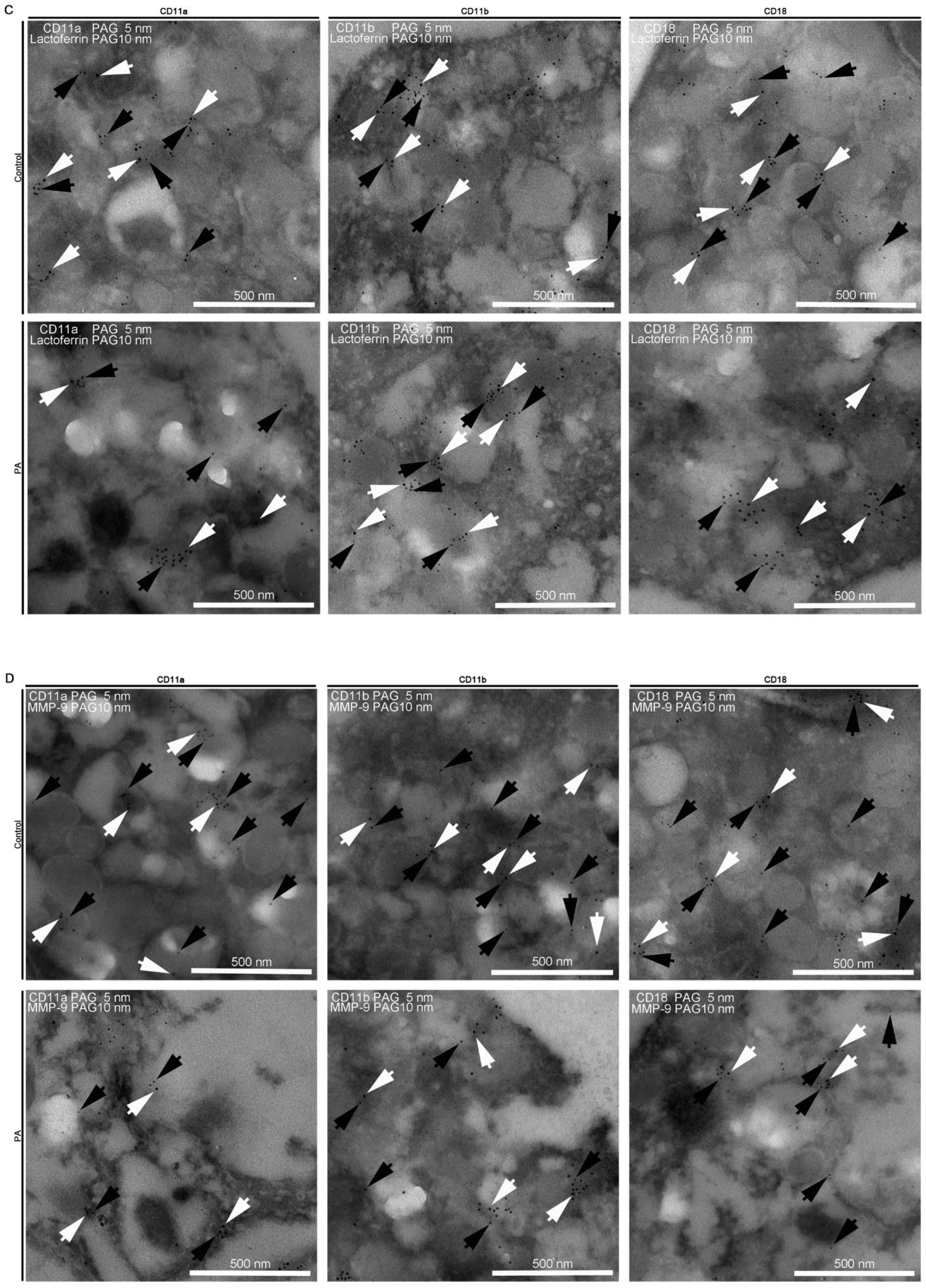
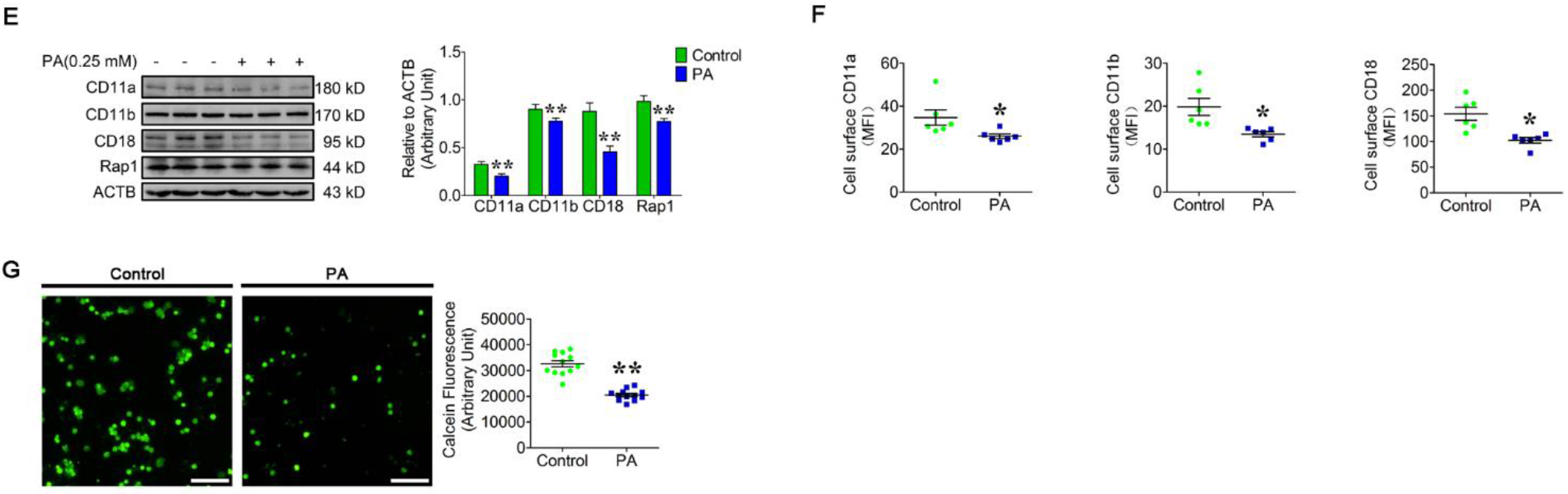
PA-induced Autophagy Decreased Neutrophil Adhesion. (A) Portion of immunogold electron micrographs of control and PA-treated neutrophils labeled with CD11a, CD11b, CD18 and Rap1 (10 nm gold grains). White arrows show gold grains on the granules, black arrows show gold grains on AVi and AVd, and v show gold grains on vacuoles. N, nucleus. p, plasma membrane. Scale bars as indicated. (B-D) Portion of immunogold electron micrographs of control and PA-treated neutrophils labeled for MPO (B), lactoferrin (C) or MMP-9 (D) (10-nm gold grains, white arrows) and then labeled for CD11a, CD11b or CD18, respectively (5-nm gold grains, black arrows). Scale bars as indicated. (E) Immunoblot for the total protein expression of CD11a, CD11b, CD18 and Rap1 in control and PA-treated neutrophils (n = 3). Data represent the mean ± s.e.m. (** P < 0.01 versus the control group; Significance calculated using two-way ANOVA). (F) Surface expression of CD11a, CD11b and CD18 on control and PA-treated neutrophils. Surface expression of CD11a, CD11b and CD18 was assessed by flow cytometry analysis (n = 6). MFI, mean fluorescence intensity. Data represent the mean ± s.e.m. (* P < 0.05 versus the control group; Significance calculated using t test). (G) Representative fluorescence micrograph images (left) and fluorescence microplate analysis (right) of control and PA-treated neutrophils adhered to HUVECs. The fluorescence intensity indicated neutrophil adhesion and was detected by a fluorescence microplate reader (n = 12). Scale bar, 400 μm. Data represent the mean ± s.e.m. (** P < 0.01 versus the control group; Significance calculated using t test).

### Hsc70-Dependent CD11a, CD11b, CD18 and Rap1 Degradation by Autophagy Reduced Neutrophil Adhesion

Ubiquitination is a prerequisite for protein degradation by autophagy^39^. We initially investigated whether CD11a, CD11b, CD18 and Rap1 could be ubiquitinated. The data showed that polyubiquitin was reciprocally coimmunoprecipitated with CD11a, CD11b, CD18 and Rap1 in neutrophils, which suggested that the IP complexes of CD11a, CD11b, CD18 and Rap1 could be polyubiquitinated (Figure 4A). The accumulation of CD11a, CD11b, CD18 and Rap1 was greatly decreased when autophagy was induced by PA, while protein levels were increased significantly when autophagy was blocked by BafA1 or CQ in PA-treated neutrophils (Figure 4B and Figure S8A). Similarly, inhibition of PA-induced autophagy increased neutrophil adhesion (Figure 4C and Figure S8B). To further confirm that PA-induced autophagy decreased neutrophil adhesion by promoting CD11a, CD11b, CD18, and Rap1 degradation, HL-60 cells were transduced with shRNAs specific for ATG5 (Figure 4D, Figure S9A, Figure S9B and Figure S9C). ATG5 knockdown attenuated PA-induced vacuolation (Figure S9A and Figure S9B), reduced the degradation of CD11a, CD11b, CD18 and Rap1 (Figure 4D and Figure S9C) and partially restored cell adhesion (Figure 4E and Figure S9D).

**Figure 4.**
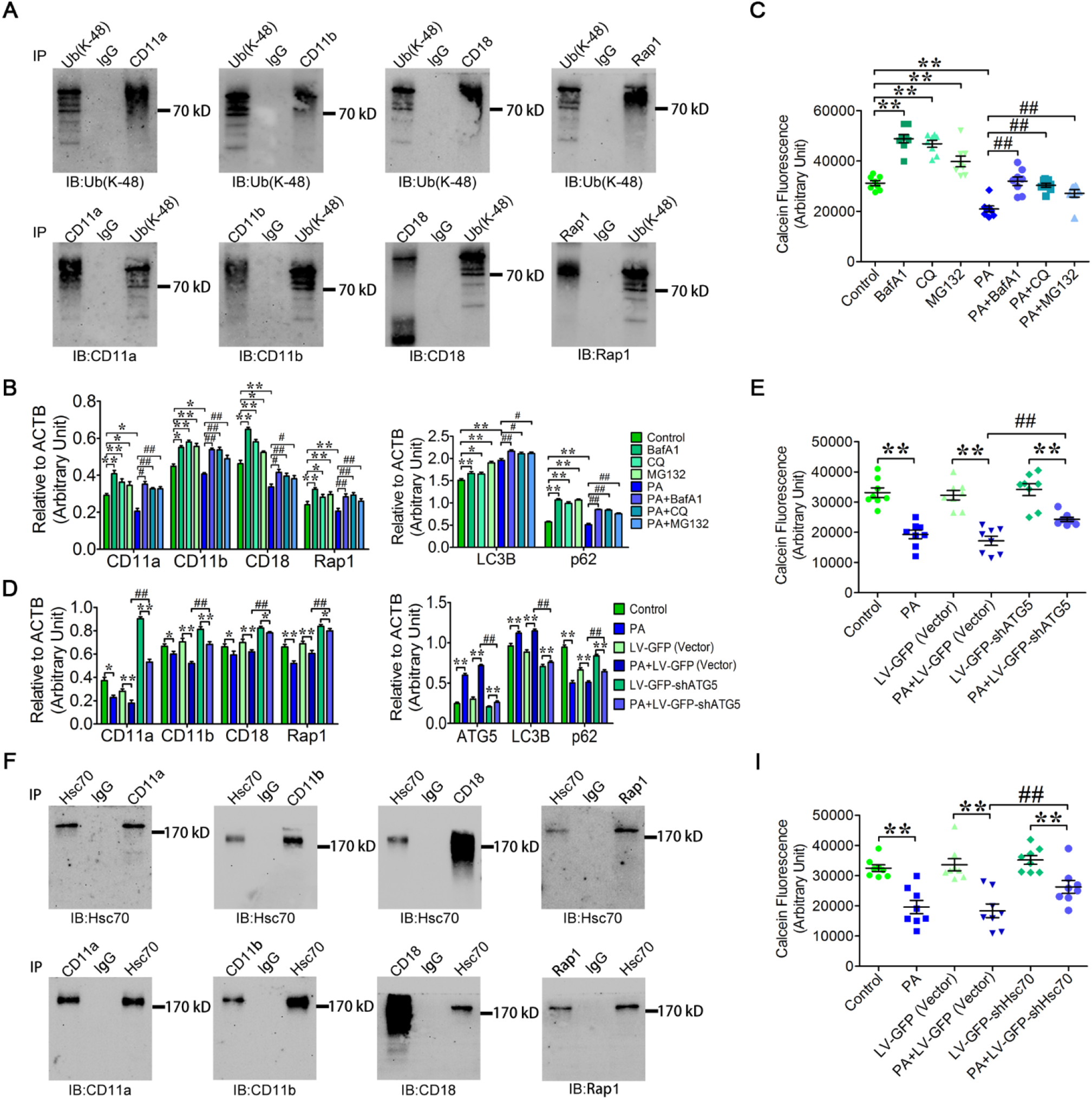
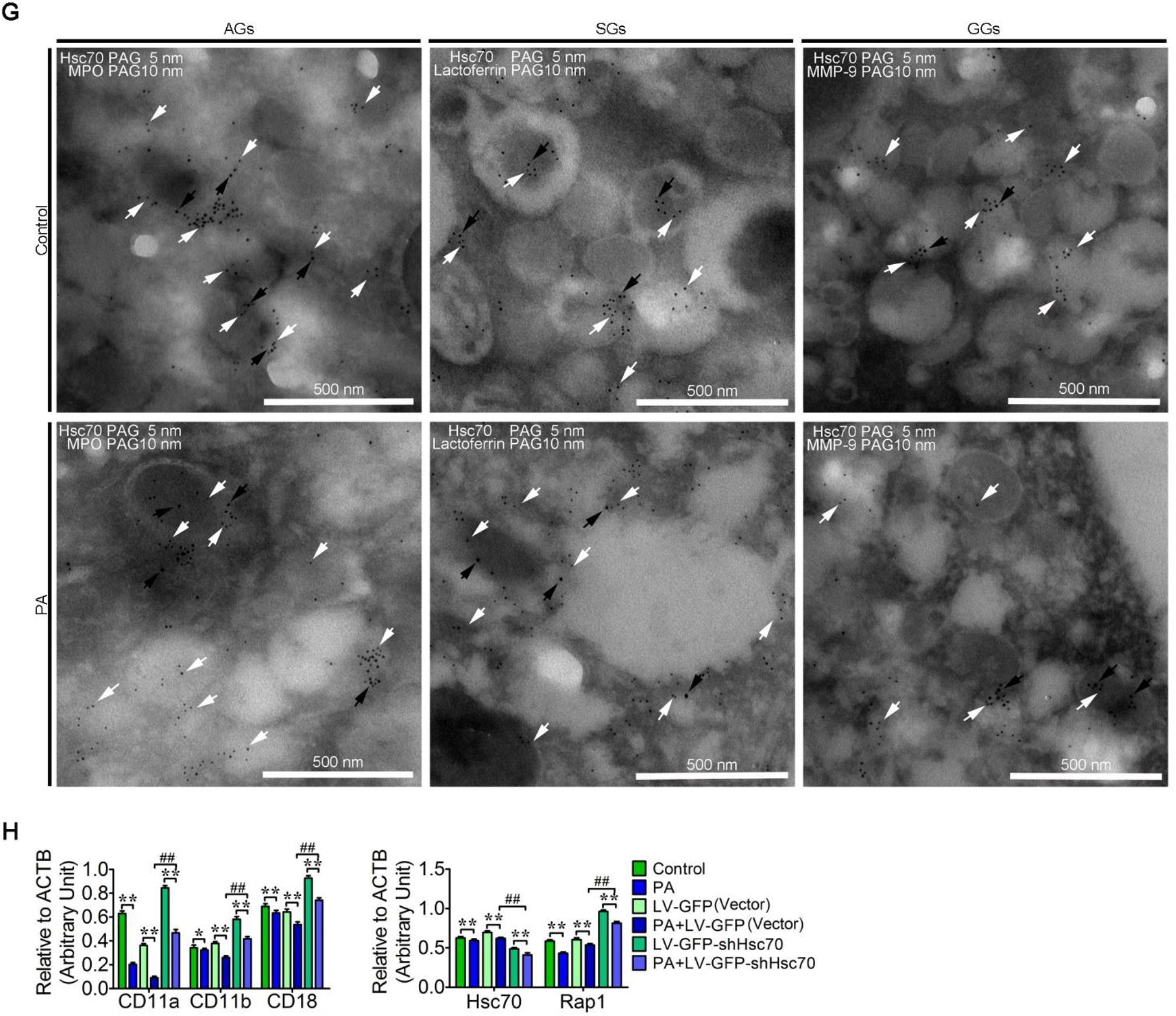
Hsc70-Dependent CD11a, CD11b, CD18 and Rap1 Degradation by Autophagy Reduced Neutrophil Adhesion. (A) The CD11a, CD11b, CD18 and Rap1 protein complexes were modified with polyubiquitin chains (Lys48). CD11a, CD11b, CD18, Rap1 and the polyubiquitinated proteins were immunoprecipitated from neutrophils and then evaluated by immunoblotting. (B-C) Inhibition of the degradation of CD11a, CD11b, CD18 and Rap1 increased neutrophil adhesion. The protein degradation was blocked by BafA1, CQ or MG132 in control and PA-treated neutrophils. Quantitative analysis of LC3B, p62, CD11a, CD11b, CD18 and Rap1 (B, n = 3) was performed, and neutrophil adhesion was detected with a fluorescence microplate reader (C, n = 8). Data represent the mean ± s.e.m. (* P < 0.05 and ** P < 0.01 versus the control group, # P < 0.05 and ## p < 0.01 versus the PA-treated group; Significance calculated using two-way ANOVA). (D-E) Knockdown of ATG5 significantly inhibited the autophagy and increased the adhesion of HL-60 cells. The HL-60 cells were infected with LV-GFP-shATG5 (to block autophagy) and LV-GFP (Vector) (negative controls). Quantitative analysis of LC3B, p62, ATG5, CD11a, CD11b, CD18 and Rap1 (D, n = 3) was performed to evaluate autophagic flux and protein accumulation. The adhesion of HL-60 cells was detected by a fluorescence microplate reader (E, n = 8). Data represent the mean ± s.e.m. (* P < 0.05 and ** P < 0.01 versus the control group, ## p < 0.01 versus the PA-treated group; Significance calculated using two-way ANOVA). (F) Reciprocal co-IP of CD11a, CD11b, CD18 and Rap1 with Hsc70. The CD11a, CD11b, CD18, Rap1 and Hsc70 protein complexes were immunoprecipitated and evaluated by immunoblotting individually. (G) Portion of immunogold electron micrograph showing the localization of Hsc70 (5 nm, white arrows) with MPO, lactoferrin or MMP-9 (10 nm, black arrows) in control and PA-treated neutrophils. Scale bars as indicated. (H-I) Knockdown of Hsc70 blocked the degradation of CD11a, CD11b, CD18 and Rap1 and increased the adhesion of HL-60 cells. The cells were infected with LV-GFP-shHsc70 and LV-GFP (Vector) (negative controls). Quantitative analysis of Hsc70, CD11a, CD11b, CD18 and Rap1 was performed (H, n = 3). Neutrophil adhesion was detected by a fluorescence microplate reader (I, n = 8). Data represent the mean ± s.e.m. (* P < 0.05 and ** P < 0.01 versus the control group, ## p < 0.01 versus the PA-treated group; Significance calculated using two-way ANOVA).

To identify the molecules involved in the degradation of the adhesion molecules and Rap1, proteins present in IP complexes of CD11a, CD11b, CD18 and Rap1 were identified by a shotgun analysis. A total of 415 proteins interacted with CD11a (Table S1); 217 proteins were identified in CD11b complexes (Table S2); 236 proteins were with CD18 (Table S3); and 399 proteins with Rap1 (Table S4). Interestingly, neither ubiquitinated peptides of CD11a, CD11b, CD18 and Rap1 nor peptides of p62 were detected. This suggested that CD11a, CD11b, CD18 and Rap1 were not directly ubiquitinated and recognized by p62. Unexpectedly, a total of 27 common proteins interacted with CD11a, CD11b, CD18 and Rap1 (Table S5), including the molecular chaperone Hsc70. Hsc70 is known to target and then deliver cytosolic proteins to lysosomes for degradation via the chaperone molecular autophagy (CMA) pathway ^40^. We confirmed the interaction as Hsc70 was reciprocally coimmunoprecipitated with CD11a, CD11b, CD18 and Rap1 (Figure 4F). In addition, immunogold electron microscopy results showed that Hsc70 gold particles were present on the granules in control neutrophils (Figure S10). However, Hsc70 immunogold signal was observed on AVi and AVd but rarely present on the vacuoles in PA-treated neutrophils (Figure S10). This suggested that Hsc70 was targeting the adhesion molecules and Rap1 for lysosomal degradation.

The three granule subunits of neutrophils are analogous to classic lysosomes as they contain LAMPs, p62 and proteolytic enzymes^41,42^. However, AGs are viewed as turnover factories of ubiquitinated protein aggregates in neutrophils^43^. As a molecular chaperone, Hsc70 can target and deliver the cytosolic proteins to lysosomes for degradation by CMA^40^. To further investigate whether Hsc70 is involved in the delivery of CD11a, CD11b, CD18 and Rap1 to the lysosome-like granules during PA-induced autophagy, colocalization analysis of Hsc70 with MPO, lactoferrin and MMP-9, as well as the morphological analysis of the three granule subunits were performed in neutrophils. Hsc70 colocalized with MPO on AGs, lactoferrin on SGs, and MMP-9 on GGs (Figure 4G). This was consistent with prior results showing Hsc70 distribution on the types of granule^44^. To confirm the role of Hsc70 in the degradation of CD11a, CD11b, CD18 and Rap1, HL-60 cells were transduced with a lentivirus that produced shRNAs specific for Hsc70. Hsc70 knockdown attenuated the PA-induced degradation of CD11a, CD11b, CD18 and Rap1 (Figure 4H and Figure S11A) and adhesion of HL-60 cells was partially restored (Figure 4I and Figure S11B). These findings indicated that CD11a, CD11b, CD18 and Rap1 were partially targeted by Hsc70, delivered to granules and degraded by autophagy following PA treatment. However, the mechanism of Hsc70 shuttles the CD11a, CD11b, CD18 and Rap1 to the three lysosomal like granules warrant further investigation.

### PA Induced Autophagy via the p-PKCα/PKD2 Pathway and Further Decreased Neutrophil Adhesion

To elucidate the mechanisms underlying the autophagy induced by PA, quantitative proteomic analysis of control and PA-treated neutrophils was performed using iTRAQ. A total of 296 differentially expressed proteins were identified, of which 46 were upregulated proteins (P/C > 1.2, P < 0.05) and 250 were downregulated proteins (P/C < 0.833, P < 0.05) (Table S6). Intriguingly, some upregulated proteins were involved in the ubiquitin-dependent autophagic catabolic processes (Accession: A0A0U1ZID9; Q15819; P15374) and in proteolysis (B4DPA4; Q4KMP7; E5RGM3). Many downregulated proteins were involved in chemotaxis (Q9BZL6; H3BMK2; P01137), endocytosis (A0A075B6N7; A0A087WXP0), polarity (F5GZG1), migration (P01137; Q9BZL6) and adhesion (B4DNT6; P05556; Q9BZL6). Notably, the iTRAQ results showed that a downstream effector of PKCα, namely, serine/threonine protein kinase D2 (PKD2, Accession: Q9BZL6), was significantly downregulated. Consistent with the iTRAQ results, immunoblotting results showed that the expression of p-PKCα and PKD2 were significantly decreased in PA-treated neutrophils (Figure 5A) and in NAFLD neutrophils (Figure 5B). These results indicated that PA inhibited the p-PKCα/PKD2 pathway. Knockout or pharmacological inhibition of PKCα dramatically increased autophagy^22,23^. To investigate whether PA induced neutrophil autophagy and vacuolation via inhibiting the p-PKCα/PKD2 pathway and further decreased neutrophil adhesion, neutrophils were treated with the p-PKCα/PKD2 inhibitor GO6983 with or without PA. PA or GO6983 significantly inhibited the p-PKCα/PKD2 pathway and upregulated the lipidation levels of LC3B, downregulated p62 accumulation (Figure 5D), and increased neutrophil vacuolation (Figure 5C). Treatments also decreased CD11a, CD11b, CD18 and Rap1 protein levels and impaired neutrophil adhesion (Figure 5E and Figure 5F). Transduction of dHL-60 cells with a PRKD2-overexpressing lentivirus attenuated the effect of PA on autophagy (Figure 5G, Figure 5H and Figure 5I), vacuolation (Figure 5G and Figure 5H), and adhesion (Figure 5J and Figure 5K), as evidenced by the significantly decreased lipidation of LC3B, increased accumulation of p62, CD11a, CD11b, CD18 and Rap1 (Figure 5I), and improved adhesion of dHL-60 cells (Figure 5J and Figure 5K). Taken together, these findings indicated that PA inhibited p-PKCα/PKD2 pathway, leading to autophagy and vacuolation, and the subsequent decreased expression of CD11a, CD11b, CD18 and Rap1 and neutrophil adhesion.

**Figure 5.**
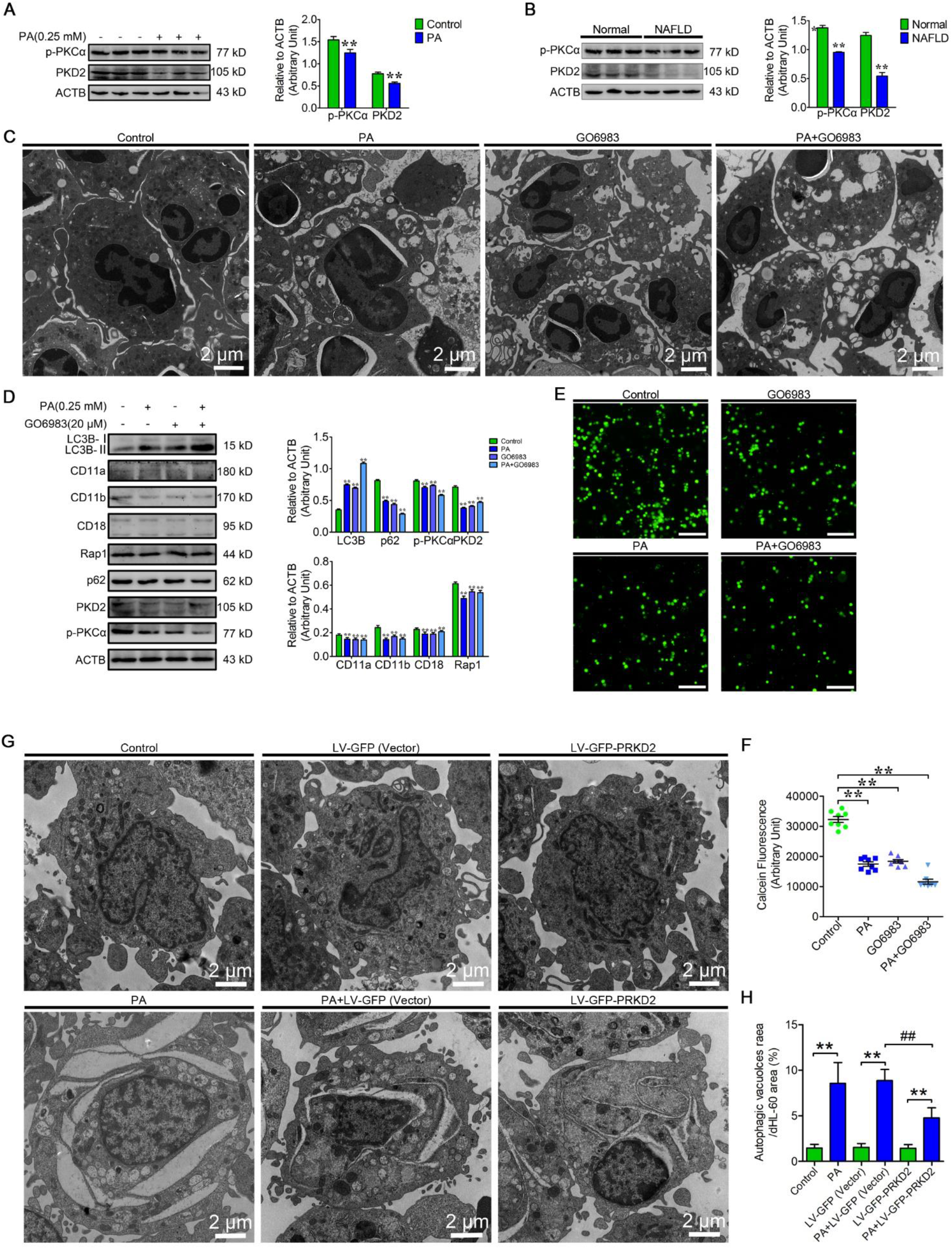
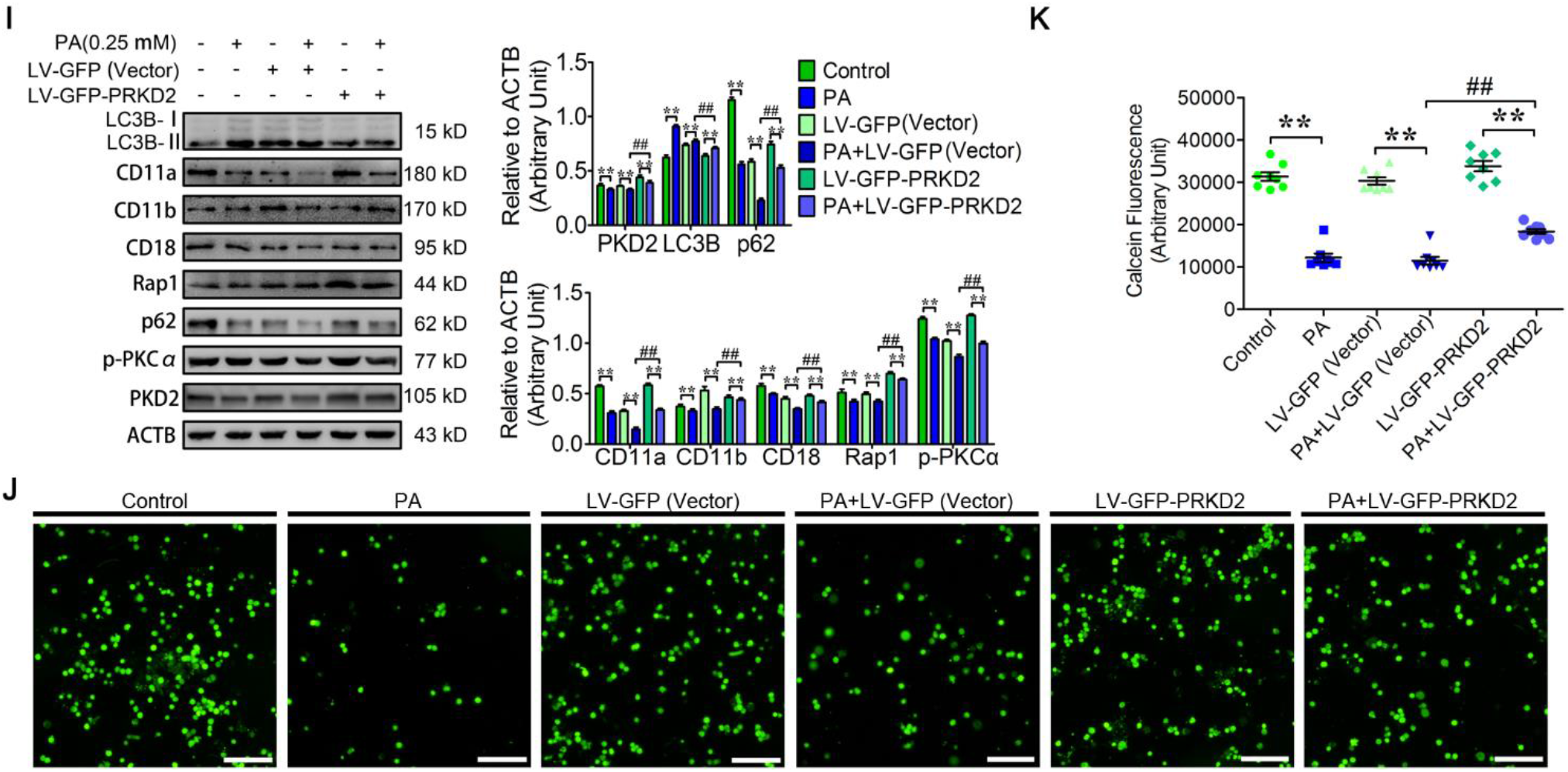
PA Induced Autophagy via the p-PKCα/PKD2 Pathway and Further Decreased Neutrophil Adhesion. (A) Immunoblot for p-PKCα and PKD2 in control and PA-treated neutrophils (n = 3). Data represent the mean ± s.e.m. (** P < 0.01 versus the control group; Significance calculated using two-way ANOVA). (B) Immunoblot for p-PKCα and PKD2 in normal and NAFLD neutrophils (n = 3). Data represent the mean ± s.e.m. (** P < 0.01 versus the control group; Significance calculated using two-way ANOVA). (C-F) The p-PKCα/PKD2 pathway is involved in the regulation of neutrophil autophagy and adhesion. Neutrophils were treated by PA, PKCα/PKD2 inhibitor GO6983 (10 μM). (C) Representative transmission electron micrographs of neutrophils treated by PA, GO6983. (D) Immunoblotting for p-PKCα, PKD2, LC3B, p62, CD11a, CD11b, CD18 and Rap1 was performed in PA or GO6983 treated neutrophils (n = 3). Data represent the mean ± s.e.m. (** P < 0.01 versus the control group, Significance calculated using two-way ANOVA). (E) Representative fluorescence micrographs of the corresponding treatments of neutrophils adhered to HUVECs. Scale bar, 400 μm. (F) Neutrophil adhesion was detected using a fluorescence microplate reader (n = 8). Data represent the mean ± s.e.m. (** P < 0.01 versus the control group; Significance calculated using one-way ANOVA). (G-K) PKD2 overexpression attenuated PA-induced autophagy and increased the adhesion of HL-60 cells. The cells were infected or not infected with LV-GFP-PRKD2 and LV-GFP (Vector) and then to induce autophagy with PA. (G) Representative transmission electron micrographs of different groups dHL-60 cells. (H) the area ratio of autophagic vacuoles to dHL-60 cells were determined (n = 6). Data represent the mean ± s.e.m. (** P < 0.01 versus the control group, ## p < 0.01 versus the PA-treated group; Significance calculated using two-way ANOVA). (I) Immunoblot for LC3B, p62, p-PKCα, PKD2, CD11a, CD11b, CD18 and Rap1 (n = 3) were performed in HL-60 cells. (J) Representative fluorescence micrographs of the corresponding treatments of HL-60 cells adhered to HUVECs. Scale bar, 400 μm. (K) Fluorescence microplate analysis of HL-60 adhesion was detected by a fluorescence microplate reader (n = 8). Data represent the mean ± s.e.m. (** P < 0.01 versus the control group, ## p < 0.01 versus the PA-treated group; Significance calculated using two-way ANOVA).

## Discussion

Integrins are required for cancer cell matrix adhesion and firm adhesion of neutrophils^45,46^. Autophagy decreases cancer cell matrix adhesion and facilitates tumor metastasis by degrading β1 integrins^47^. However, it is unknown whether autophagy decreases the firm adhesion of neutrophils by degrading β2 integrins. In metabolic diseases, neutrophils are exposed to abnormal metabolite levels, such as high blood levels of fatty acids, which exhibit lipotoxicity and can impair neutrophil immune function. In this study, we found that the three neutrophil granule types, namely, AGs, SGs and GGs, could be engulfed by autophagosomes for degradation in NAFLD neutrophils. Furthermore, CD11a, CD11b, CD18 and Rap1 in the neutrophils were targeted by Hsc70 and degraded via autophagy. Consequently, neutrophil adhesion was significantly decreased. Notably, we found that PA inhibited the p-PKCα/PKD2 pathway to induce autophagy. In neutrophils, autophagic vacuoles exhibit morphological diversity, and the classification of these vacuoles is not well standardized. Many appellations, such as phagocytic vacuole^48^, glycogen autophagosome^49^ and vacuole^50^, have been used to describe neutrophil autophagic vacuoles. We first divided the neutrophil autophagic vacuoles according to four consecutive stages, namely, AVi, AVd, glycogen vacuole and vacuole, depending on the degree of degradation of the engulfed granules or other cytosolic cargoes. The autophagy receptorp62 mediates the degradation of the damaged mitochondria in energy cells^51^. We found that most AGs, SGs and GGs colocalized with p62 and MPO, lactoferrin and MMP-9, respectively, were located on the AVi and AVd of neutrophil autophagic vacuoles. Little to no signal was observed on the glycogen vacuole and vacuole stages, which indicated that p62 might deliver the damaged AGs, SGs and GGs to lysosomes for degradation via autophagy. Interestingly, granules are also considered as the lysosomes of neutrophils^52,53^. Damaged lysosomes can be eliminated through autophagy^54^. Moreover, lysosomes can fusion with autophagosomes, which process further damaged lysosomes, as well the lysosomal proteins are released into the cytoplasm. Degranulation is the process of regulated exocytosis of these lysosome-like granules. Autophagy deficiency inhibits degranulation^55^. Whether the fusion of the granules with autophagosomes plays a role in the regulation of degranulation warrant further investigation.

This continuous autophagic flux contributed to neutrophil vacuolation. As mentioned above, the immunity of patients with severe vacuolated neutrophils is decreased^34^. Interestingly, we found that autophagic vacuoles also existed in NAFLD neutrophils, hinting at a reduced immunity of NAFLD patients. Interestingly, CD11a, CD11b and CD18 were all observed on the AGs, SGs and GGs. These β2 integrins protein levels were lowered concomitantly with the number of the three granule types. The decreased protein level of the adhesion molecules and the upstream signaling molecule Rap1 was dependent on autophagy, and impaired neutrophil adhesion. Rap1, a β2 integrin activity regulator^8^. These findings suggested that autophagy decreases neutrophil adhesion by both degrading CD11a, CD11b and CD18 and reducing the activity of these proteins by facilitating Rap1 degradation.

Although we confirmed that the CD11a, CD11b, CD18 and Rap1 could be degraded through autophagy and autophagy plays an important role in neutrophil adhesion, we also showed that the proteasome inhibitor MG132 rescued CD11a, CD11b, CD18 and Rap1 protein levels as well as a restored cell adhesion. However, the inhibition of autophagy had a more profound effect on adhesion than the inhibition of the proteasome pathway, supporting a more important role of autophagy in the regulation of integrins in neutrophils adhesion. Ubiquitination is a prerequisite for autophagy-dependent protein degradation^56^. Multiple pathways are involved in the direct^56^ and indirect ubiquitination ^57^ of cell surface proteins. To investigate the ubiquitination of CD11a, CD11b, CD18 and Rap1, four IP complexes were enriched from neutrophils and identified by a shotgun proteomic approach. Surprisingly, no ubiquitinated peptides of CD11a, CD11b, CD18 and Rap1 were detected, possibly because the ubiquitinated protein lev els were too low or the proteins could not be ubiquitinated directly. Moreover, polyubiquitin proteins (P0CG48, Table S1 and Table S2 and Table S4) and E3 ubiquitin-protein ligases (Q76N89, O76064, Q86UK7, Table S1; Q86Y13, H7C3Z1 Table S3; Q9NQC1, A0A096LP02, Q96T88, Table S4) were identified in the IP complexes, suggesting that these four proteins might form polyubiquitin-protein conjugates and might be degraded by p62-dependent autophagy. The α5β1 integrin could be degraded in a ligand (fibronectin)-dependent manner^58^. However, no β2 integrin ligands (FGA, FGB and FGG) and p62 were observed in the mass spectrometry results, which indicated that the degradation of CD11a, CD11b, CD18 and Rap1 was not dependent on ligands or p62 in neutrophils. Notably, Hsc70 (P11142) was identified from four IP complexes and could be reciprocally coimmunoprecipitated with CD11a, CD11b, CD18 and Rap1. Furthermore, CD11b, CD18, and AG, SG, and GG marker proteins (MPO: P05164, lactoferrin: P02788 and MMP-9: P14780, respectively) were also observed in the IP complexes of Hsc70 by the shotgun approach (Table S7). Coincidentally, MPO, lactoferrin and MMP-9 were also identified in the IP complexes of CD11a, CD11b, CD18 and Rap1. In addition, Hsc70 colocalized with MPO, lactoferrin and MMP-9 on AGs, SGs and GGs, respectively. These results suggested that CD11a, CD11b, CD18 and Rap1 were delivered by Hsc70 to lysosomes for degradation. However, the special motif (KFERQ) was not found in peptides of the CD11a, CD11b, CD18 and Rap1. Hsc70 might a interact with a partner protein of the adhesion molecules with the motif and delivered them to the lysosome for degradation. Notably, Hsc70 knockdown attenuated autophagy-mediated degradation of CD11a, CD11b, CD18 and Rap1, thereby increasing the adhesion of HL-60 cells. Taken together, the results showed that CD11a, CD11b, CD18 and Rap1 could be degraded via autophagy.

Our data demonstrated that autophagy induced by PA decreased the adhesion of PA-treated and NAFLD neutrophils. However, the underlying mechanism was unclear. Interestingly, using quantitative proteomic analysis, we found that a downstream target of PKCα, namely, PKD2^59^, a key regulatory protein of autophagy^60^, was significantly downregulated. PKCα is a negative regulator of autophagy in neuroepithelial cells^23^. Our data showed that PKCα/PKD2 was indeed inhibited in PA-treated and NAFLD neutrophils. Moreover, pharmacological inhibition of PKCα/PKD2 by GO6983 strongly induced neutrophil autophagy, vacuolation and decreased neutrophil adhesion, while PKD2 overexpression significantly attenuated the PA-induced autophagy, vacuolation and decrease in adhesion. These results indicated that PKCα/PKD2 pathway was involved in PA-induced autophagy and then caused neutrophil vacuolation and a decrease in adhesion. A previous study indicated that PKC inhibitors dramatically induced autophagy^22^, which further support our conclusion.

Collectively, this study reveals that PA inhibited the p-PKCα/PKD2 pathway to induce autophagy (in vitro and *ex vivo*), which caused neutrophil vacuolation, promoted the degradation of CD11a, CD11b, CD18 and Rap1 and further decreased neutrophil adhesion, thereby impairing neutrophil immunity. Notably, we found that the three neutrophil granule subunits, namely, AGs, SGs and GGs, were degraded by autophagy. This phenomenon might be termed “granulophagy” (a combination of “granule” and “autophagy”). In addition to adhesion-associated proteins, proteins associated with endocytosis, phagocytosis, phagosomes, chemotaxis, microbicidal substances, cytoskeleton remodeling, etc. are also present on AGs, SGs, and GGs^2^, and these proteins might be degraded by autophagy. It still remains to be determined if, besides adhesion, the ability of neutrophils to migrate and to kill microbes is also affected by PA or is impaired in NAFLD PMNs. Autophagy might be an immune switch for neutrophils that control the above biological functions. Understanding the mechanisms of autophagy-induced degradation of intracellular immune-associated proteins is fundamental to the identification of new therapeutic strategies against metabolic disease-induced innate immune deficiency. Neutrophil vacuolation, an indicator of the immune status of patient^34^, is associated with autophagy resulting from changes in blood constituents. Therefore, further research is needed to investigate the effects of the metabolic disorders on the autophagy and vacuolation of neutrophils, which will provide a new therapeutic strategy to improve the immune deficiency resulting from the above diseases.

## Acknowledgments

This work was supported by the National Key Research and Development Program (Beijing, China; grant no. 2016YFD0501206), the National Natural Science Foundation of China (Beijing, China; grant no. 31372494, 31672621, and 31772810) and the Jilin Natural Science Foundation (Changchun, China; grant no. 20170101148JC) and the International Postdoctoral Exchange Fellowship Program (20190057). We thank Dr. Fabien Loison for assistance with editing and grammar correction.

## Authorship Contributions

Z.C.P., A.Y.H designed the study, performed experiments, analyzed most data, and wrote the manuscript; H.Y.W. contributed to the Normal and NAFLD samples and edited the manuscript; Y.C.Y., B.C.F., X.L.D., Y.F.L., Y.W.Z. performed experiments, analyzed the data and assisted in generating LV-GFP-shATG5, LV-GFP-shAp62, LV-GFP-shHsc70 and LV-GFP-PRKD2 HL-60 cells, and contributed to writing the manuscript; X.B.L., Z.W., G.W.L., X.W.L. jointly directed this work, including designed, analyzed, supervised overall project, and co-wrote the manuscript, with input from all authors.

---

**Figure S1.**
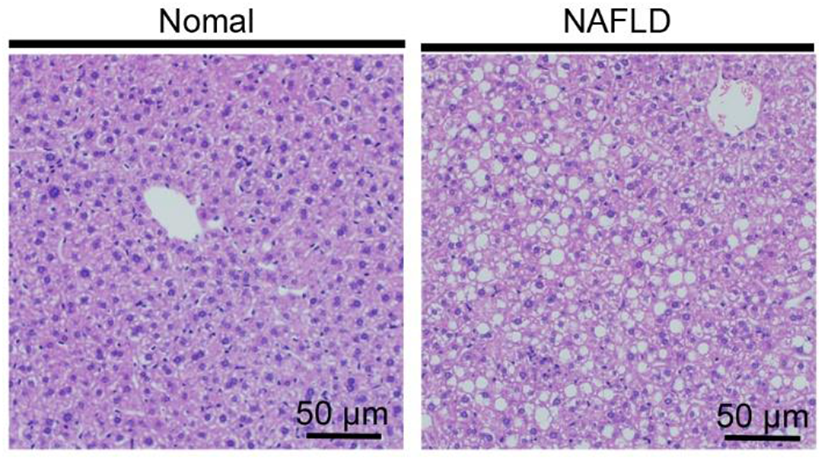
Representative images of HE-stained liver sections from normal individuals and NAFLD patients. Scale bars as indicated.

**Figure S2.**
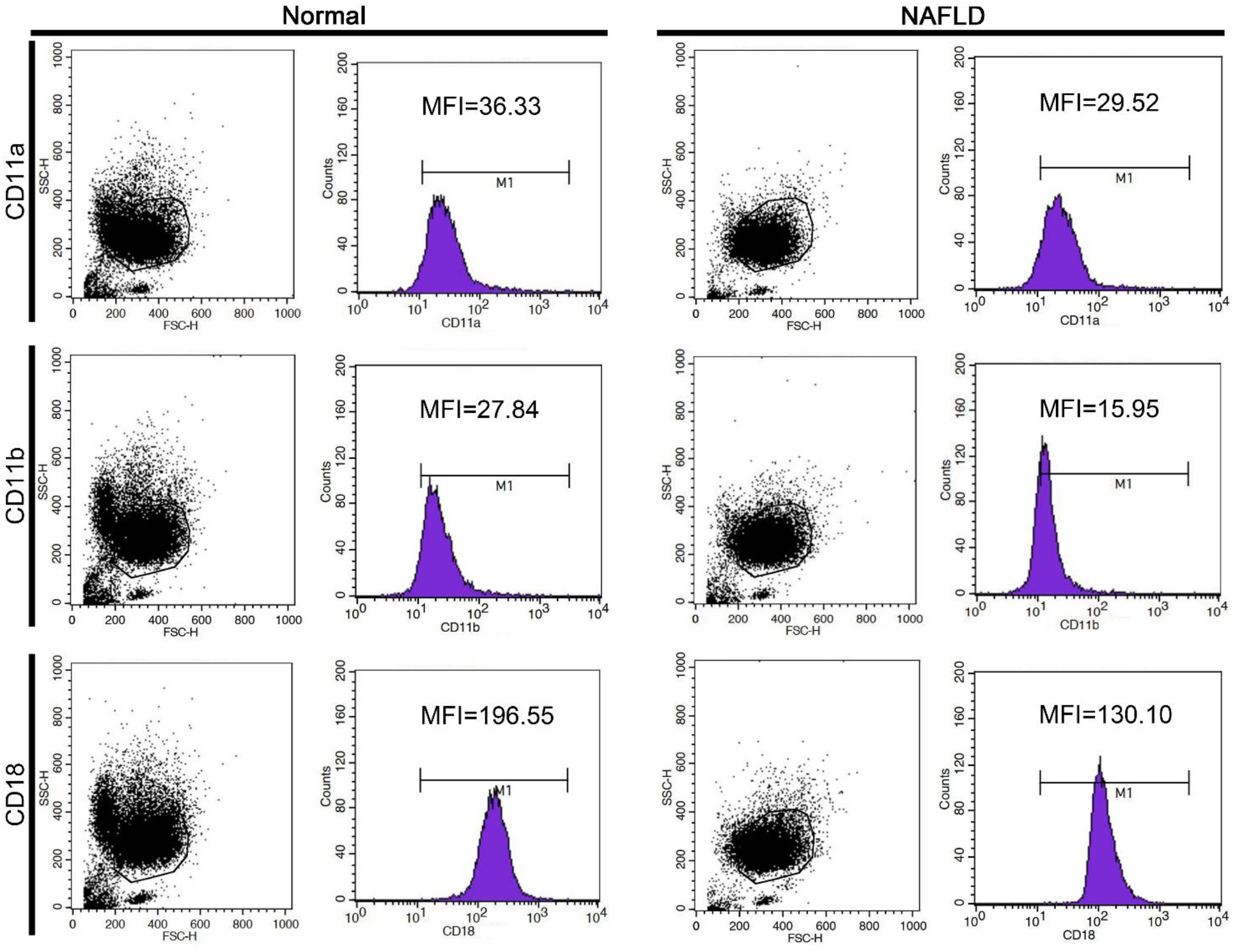
Flow cytometry images of the surface expression of CD11a, CD11b and CD18 on normal and NAFLD neutrophils.

**Figure S3.**
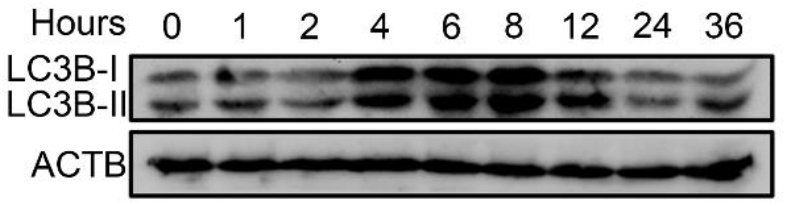
Immunoblot for the lipidation of LC3B in PA (0.25 mM)-treated neutrophils.

**Figure S4.**
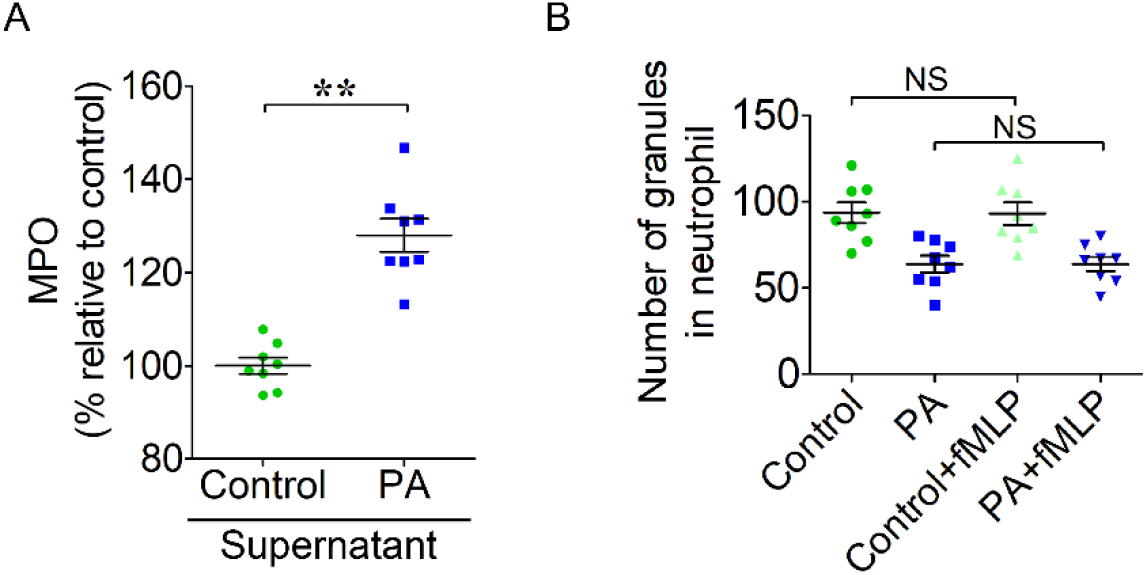
(A) ELISA results of MPO in PA-treated neutrophil supernatant. The supernatant of control and PA-treated neutrophils was collected, the content of MPO was detected with the ELISA kit. Data represent the mean ± s.e.m. (** P < 0.01 versus the control group; Significance calculated using t test). (B) The number of granules in normal and PA-treated neutrophils with or without using fMLP to induce degranulation. Control and PA-treated neutrophils were cultured for 6h and then stimulated with 1 μM fMLP for 30 min to induce degranulation and then were collected to perform the granule quantification, without fMLP treatment groups as control. Data represent the mean ± s.e.m. (NS means no different versus the not stimulated groups; Significance calculated using one-way ANOVA).

**Figure S5.**
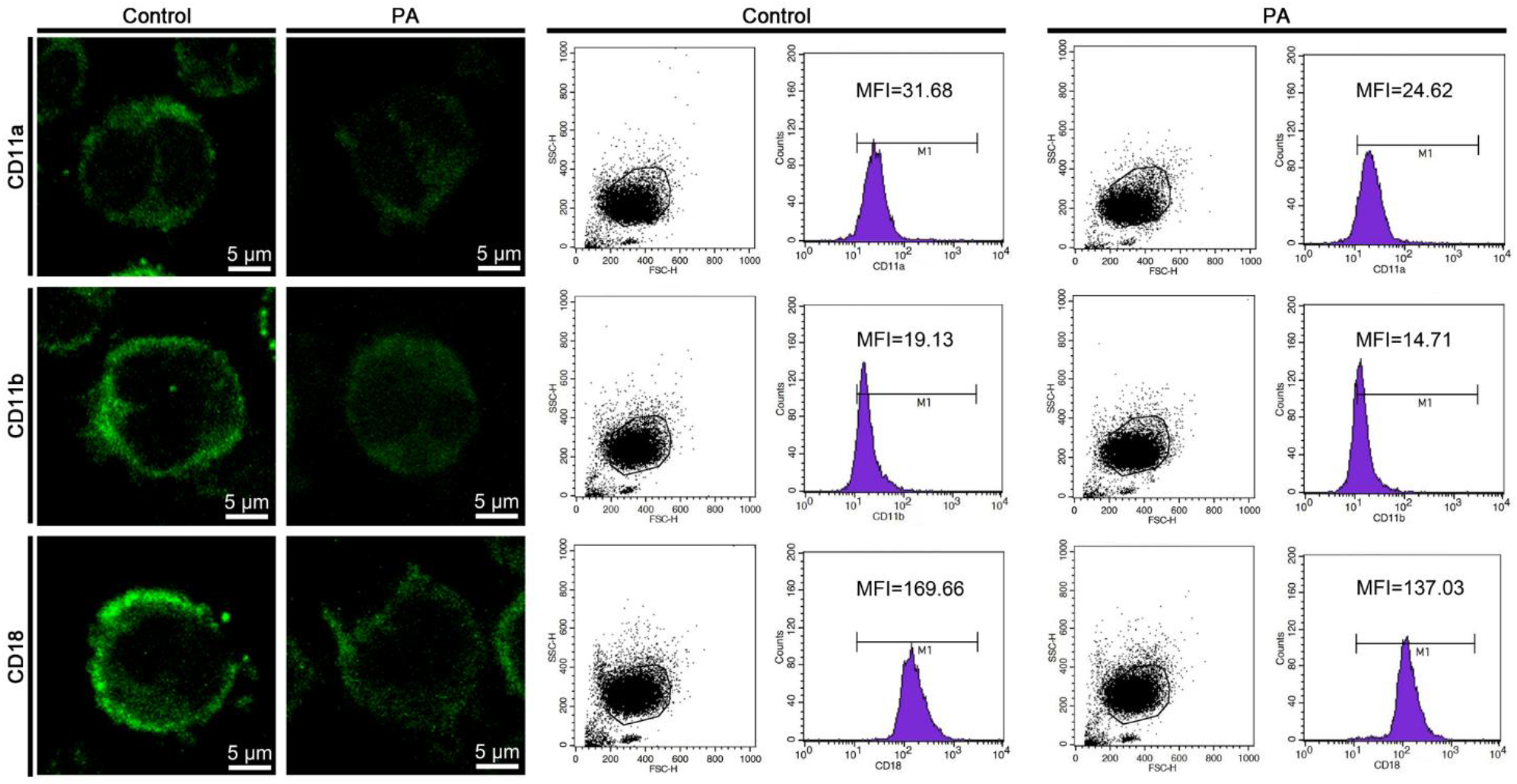
Representative immunofluorescence images (left) and flow cytometry images (right) of the surface expression of CD11a, CD11b and CD18 on control and PA-treated neutrophils. Scale bars as indicated.

**Figure S6.**
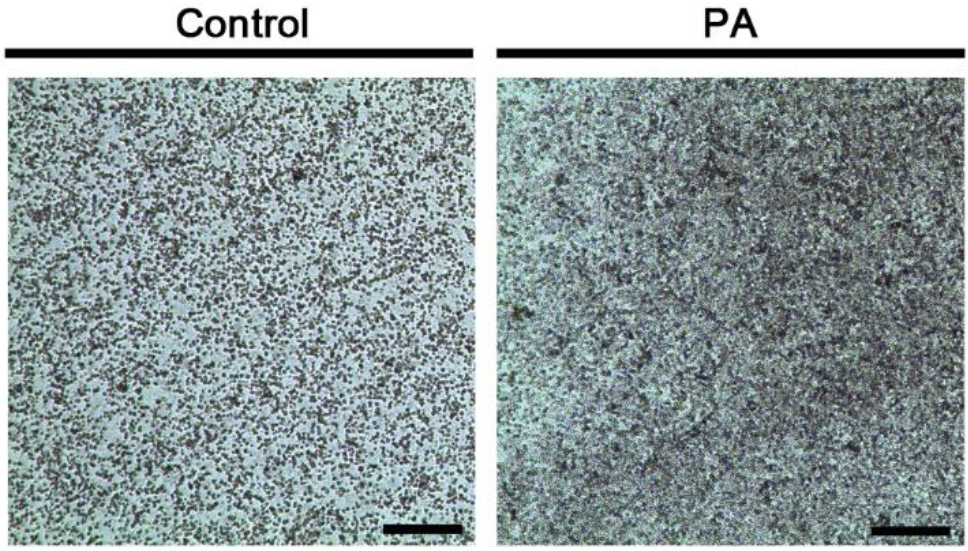
Representative light microscopy images of control and PA-treated neutrophils. Scale bar, 2 mm.

**Figure S7.**
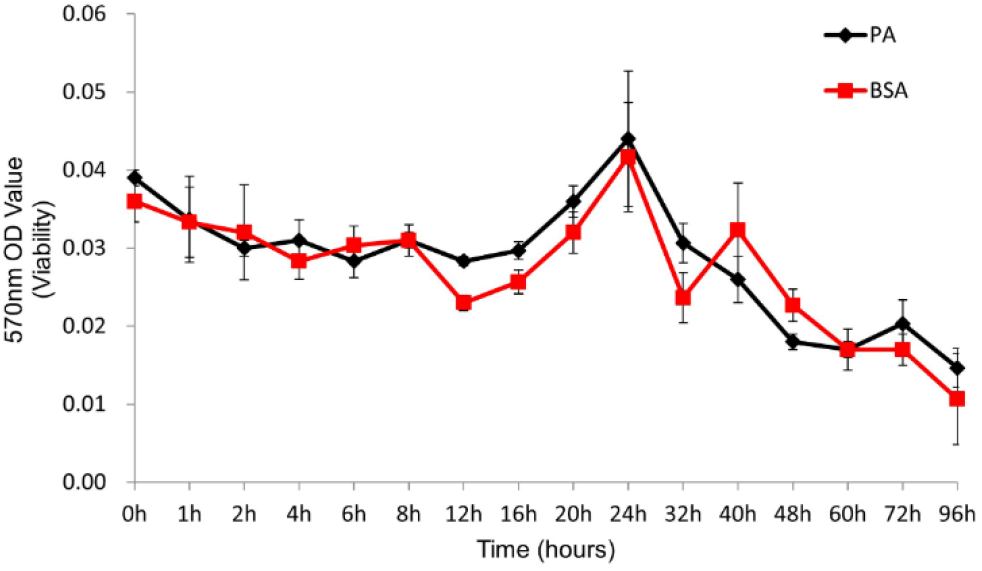
The viability of BSA control and PA (0.25 mM)-treated neutrophils was assessed using the CCK-8 cytotoxicity assay.

**Figure S8.**
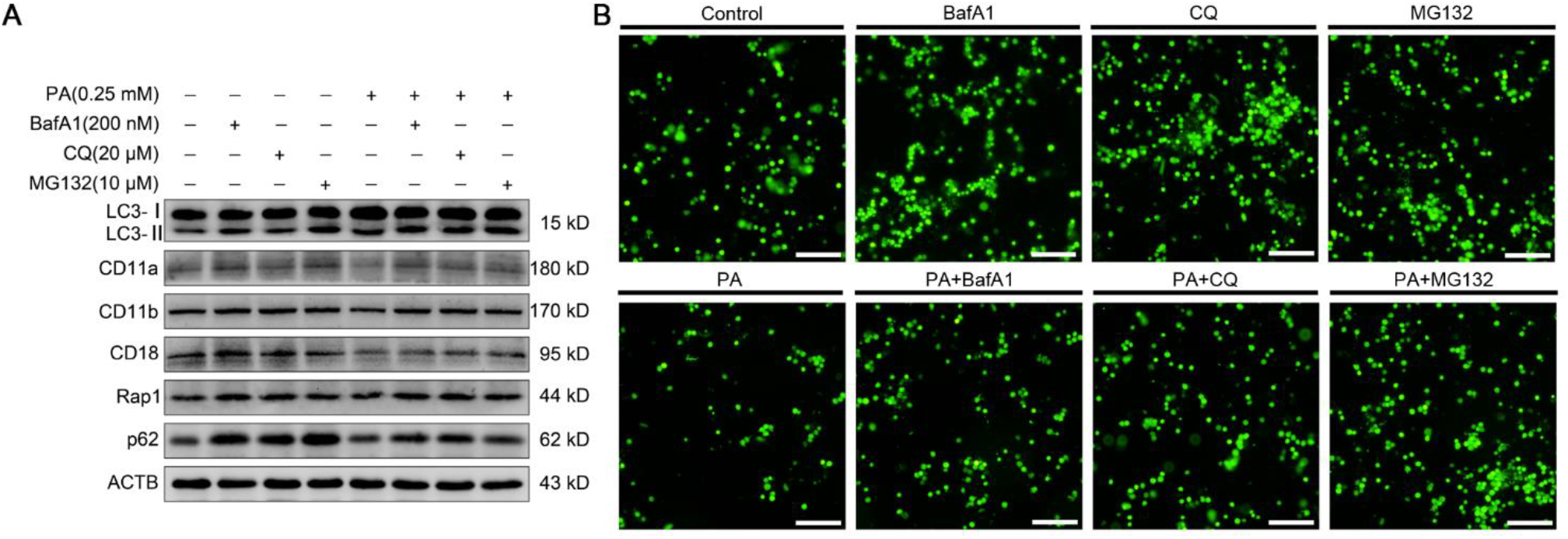
(A) Immunoblot for LC3B, p62, CD11a, CD11b, CD18 and Rap1 (A) in control and PA-treated neutrophils (treated or not treated with BafA1, CQ or MG132). (B) Representative fluorescence micrographs of control and PA-treated neutrophils (treated or not treated with BafA1, CQ or MG132) adhered to HUVECs. Scale bar, 400 μm.

**Figure S9.**
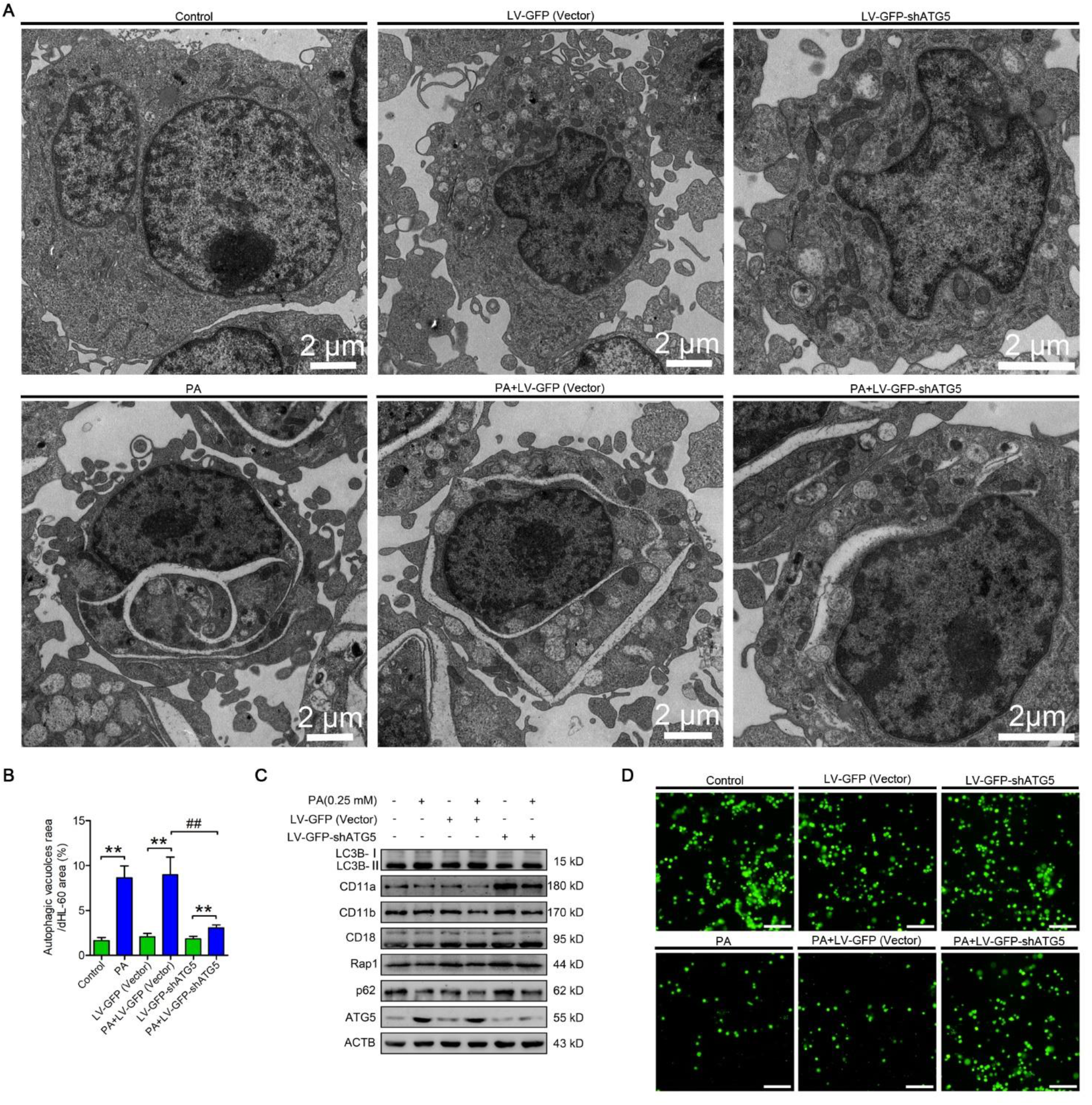
(A) Representative transmission electron micrographs of control and PA-treated HL-60 cells (infected or not infected with LV-GFP-shATG5 or empty lentivectors). Scale bars as indicated. (B) the area ratio of autophagic vacuoles to dHL-60 cells were determined (n = 6). Data represent the mean ± s.e.m. (** P < 0.01 versus the control group, ## p < 0.01 versus the PA-treated group; Significance calculated using two-way ANOVA). (C) Immunoblot for LC3B, p62, ATG5, CD11a, CD11b, CD18 and Rap1 in control and PA-treated HL-60 cells (infected or not infected with LV-GFP-shATG5 or empty lentivectors). (D) Representative fluorescence micrographs of control and PA-treated differentiated HL-60 cells (infected or not infected with LV-GFP-shATG5 or empty lentivectors) adhered to HUVECs. Scale bar, 400 μm.

**Figure S10.**
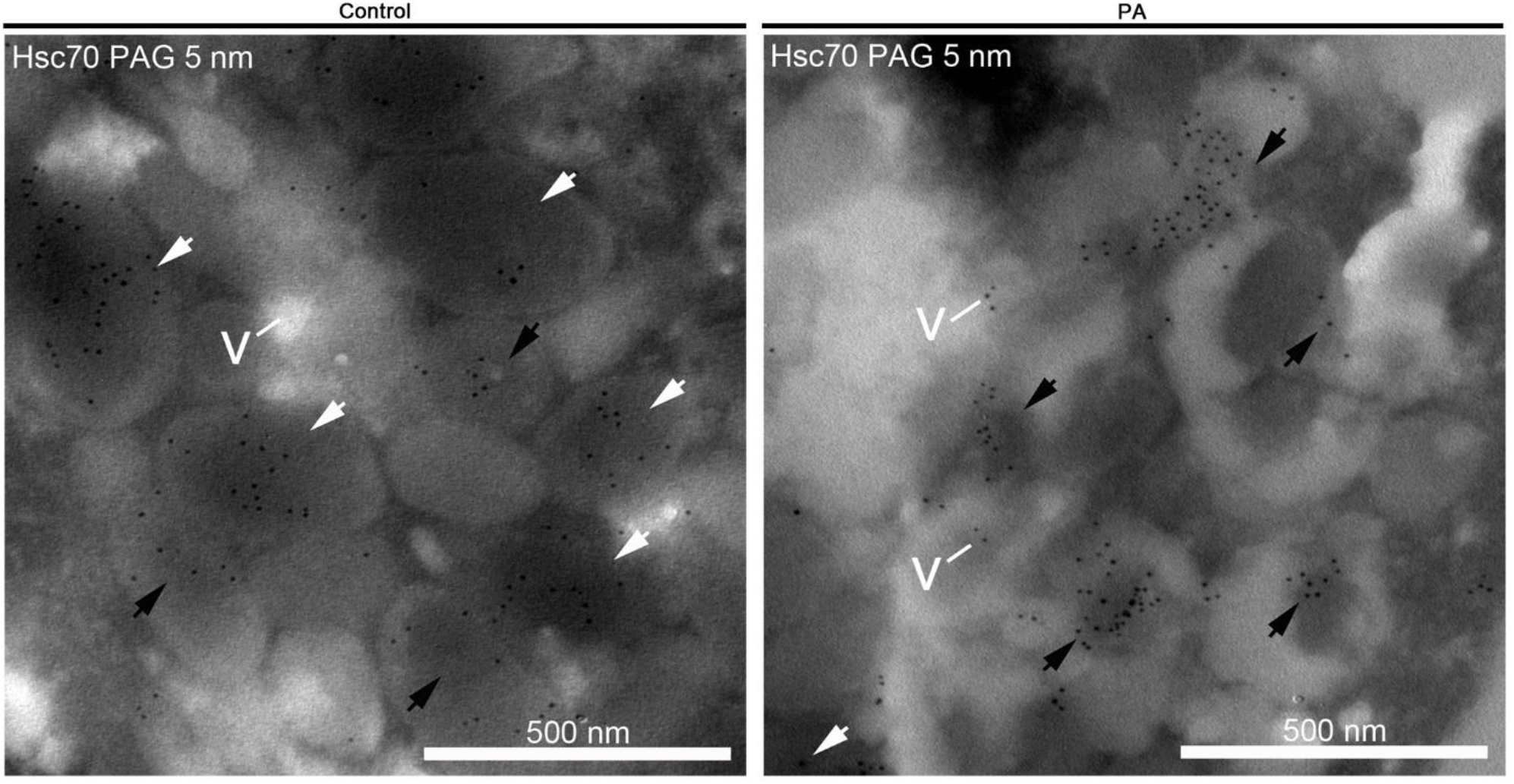
Immunogold electron micrographs showing the localization of Hsc70 in control and PA-treated neutrophils. White arrows, granules. Black arrows, AVi and AVd. v, vacuoles. Scale bars as indicated.

**Figure S11.**
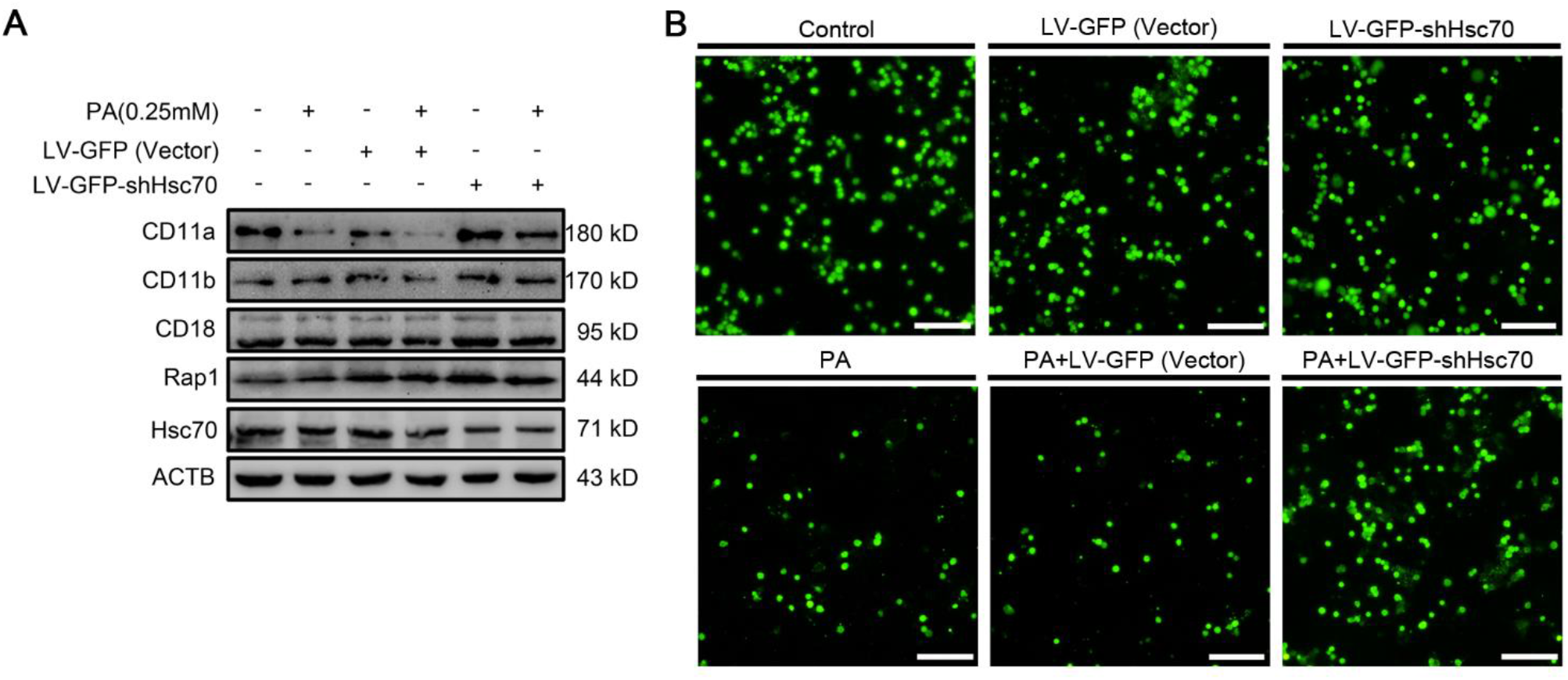
(A) Immunoblotting for Hsc70, CD11a, CD11b, CD18 and Rap1 was performed in control and PA-treated HL-60 cells (infected or not infected with LV-GFP-shHsc70 or empty lentivectors). (B) Representative fluorescence micrographs of control and PA-treated HL-60 cells (infected or not infected with LV-GFP-shHsc70 or empty lentivectors) adhered to HUVECs. Scale bar, 400 μm.

**Table S1.** The list of proteins interacting with CD11a identified by the shotgun

**Table S2.** The list of proteins interacting with CD11b identified by the shotgun

**Table S3.** The list of proteins interacting with CD18 identified by the shotgun

**Table S4.** The list of proteins interacting with Rap1 identified by the shotgun

**Table S5.** The list of common proteins interacting with CD11a, CD11b, CD18 and Rap1 identified by the shotgun

**Table S6.** The list of differentially expressed proteins between PA treatment group and control group by iTRAQ

**Table S7.** The list of proteins interacting with Hsc70 identified by the shotgun

## References

1. Németh T, Sperandio M, Mócsai A. Neutrophils as emerging therapeutic targets. Nature Reviews Drug Discovery. 2020:1–23.

2. Lominadze G, Powell DW, Luerman GC, Link AJ, Ward RA, Mcleish KR. Proteomic analysis of human neutrophil granules. Molecular & Cellular Proteomics Mcp. 2005;4(10):1503.

3. Zhang L. The α M β 2 integrin and its role in neutrophil function. Cell research. 1999;9(3):171–178.

4. Bainton DF, Miller LJ, Kishimoto TK, Springer TA. Leukocyte adhesion receptors are stored in peroxidase-negative granules of human neutrophils. Journal of Experimental Medicine. 1987;166(6):1641–1653.

5. Pellinen T, Ivaska J. Integrin traffic. Journal of Cell Science. 2006;119(Pt 18):3723–3731.

6. Williams MR, Azcutia V, Newton G, Alcaide P, Luscinskas FW. Emerging mechanisms of neutrophil recruitment across endothelium. Trends in Immunology. 2011;32(10):452–460.

7. Evans R, Patzak I, Svensson L, et al. Integrins in immunity. Journal of Cell Science. 2009;122(2):215–225.

8. Bos JL, De BK, Enserink J, et al. The role of Rap1 in integrin-mediated cell adhesion. Biochemical Society Transactions. 2003;31(1):83–86.

9. Kinashi T, Aker M, Sokolovskyeisenberg M, et al. LAD-III, a leukocyte adhesion deficiency syndrome associated with defective Rap1 activation and impaired stabilization of integrin bonds. Blood. 2004;103(3):1033–1036.

10. Dimanche MT, Le DF, Fischer A, Arnaout MA, Griscelli C, Lisowskagrospierre B. LFA-1 beta-chain synthesis and degradation in patients with leukocyte-adhesive proteins deficiency. European Journal of Immunology. 1987;17(3):417–419.

11. Abeliovich H. Guidelines for the use and interpretation of assays for monitoring autophagy. Autophagy. 2012.

12. Mihalache CC, Yousefi S, Conus S, Villiger PM, Schneider EM, Simon HU. Inflammation-associated autophagy-related programmed necrotic death of human neutrophils characterized by organelle fusion events. Journal of Immunology. 2011;186(11):6532.

13. Borregaard N, Cowland JB. Granules of the human neutrophilic polymorphonuclear leukocyte. Blood. 1997;89(10):3503–3521.

14. Jin M, Liu X, Klionsky DJ. SnapShot: Selective Autophagy. Cell. 2013;152(1-2):368–368.e362.

15. Raiborg C, Stenmark H. The ESCRT machinery in endosomal sorting of ubiquitylated membrane proteins. Nature. 2009;458(7237):445–452.

16. Chua CEL, Gan BQ, Tang BL. Involvement of members of the Rab family and related small GTPases in autophagosome formation and maturation. Cellular & Molecular Life Sciences. 2011;68(20):3349–3358.

17. Eskelinen EL. Maturation of autophagic vacuoles in Mammalian cells. Autophagy. 2005;1(1):1–10.

18. Sahu R, Kaushik S, Clement CC, et al. Microautophagy of cytosolic proteins by late endosomes. Developmental Cell. 2011;20(1):131–139.

19. Roberts, Marnie, Barry, et al. PDGF-regulated rab4-dependent recycling of αvβ3 integrin from early endosomes is necessary for cell adhesion and spreading. Current Biology. 2001;11(18):1392–1402.

20. Ng T, Shima D, Squire A, et al. PKCα regulates β1 integrin-dependent cell motility through association and control of integrin traffic. Embo Journal. 2014;18(14):3909–3923.

21. Tan SH, Shui G, Zhou J, et al. Induction of autophagy by palmitic acid via protein kinase C-mediated signaling pathway independent of mTOR (mammalian target of rapamycin). Journal of Biological Chemistry. 2012;287(18):14364–14376.

22. Jiang H, Cheng D, Liu W, Peng J, Feng J. Protein kinase C inhibits autophagy and phosphorylates LC3. Biochemical & Biophysical Research Communications. 2010;395(4):471–476.

23. Wang F, Xu C, Reece EA, et al. Protein kinase C-alpha suppresses autophagy and induces neural tube defects via miR-129-2 in diabetic pregnancy. Nature Communications. 2017;8:15182.

24. Wegner CS, Malerød L, Pedersen NM, et al. Erratum to: Ultrastructural characterization of giant endosomes induced by GTPase-deficient Rab5. Histochemistry & Cell Biology. 2010;133(1):57.

25. Kjeldsen L, Bainton DF, Sengelov H, Borregaard N. Structural and functional heterogeneity among peroxidase-negative granules in human neutrophils: identification of a distinct gelatinase-containing granule subset by combined immunocytochemistry and subcellular fractionation. 1993.

26. Kjeldsen L, Sengelov H, Lollike K, Nielsen MH, Borregaard N. Isolation and characterization of gelatinase granules from human neutrophils. 1994

27. Bainton DF, Ullyot JL, Farquhar MG. The development of neutrophilic polymorphonuclear leukocytes in human bone marrow. Journal of Experimental Medicine. 1971;134(4):907–934.

28. Cramer E, Pryzwansky KB, Villeval JL, Testa U, Breton-Gorius J. Ultrastructural localization of lactoferrin and myeloperoxidase in human neutrophils by immunogold. Blood. 1985;65(2):423–432.

29. Kjeldsen L, Bainton DF, Sengeløv H, Borregaard N. Structural and functional heterogeneity among peroxidase-negative granules in human neutrophils: identification of a distinct gelatinase-containing granule subset by combined immunocytochemistry and subcellular fractionation. Blood. 1993;82(10):3183.

30. Kjeldsen L, Sengeløv H, Lollike K, Nielsen MH, Borregaard N. Isolation and characterization of gelatinase granules from human neutrophils. Blood. 1994;83(6):1640.

31. Donghua G, Qinghe Z, Hong Z, Dongbo S. Proteomic analysis of membrane proteins of vero cells: exploration of potential proteins responsible for virus entry. Dna & Cell Biology. 2014;33(1):20–28.

32. Léger T, Garcia C, Ounissi M, Lelandais G, Camadro JM. The metacaspase Mca1p has a dual role in farnesol-induced apoptosis in Candida albicans. Molecular & Cellular Proteomics Mcp. 2015;14(1):93.

33. Yi Z, Hong X, Hao C, et al. Proteomic analysis of solid pseudopapillary tumor of the pancreas reveals dysfunction of the endoplasmic reticulum protein processing pathway. Molecular & Cellular Proteomics. 2014;13(10):2593–2603.

34. Grange MJ, Brivet F, Boumier P, Tchernia G. [Diagnostic and prognostic values of vacuolated polymorphonuclear neutrophils (author’s transl)]. La Nouvelle Presse Medicale. 1980;9(35):2553.

35. Chetty-Raju N, Cook R, Erber WN. Vacuolated neutrophils in ethanol toxicity. British Journal of Haematology. 2004;127(5):478.

36. Zieve PD, Haghshenass M, Blanks M, Krevans JR. Vacuolization of the neutrophil. An aid in the diagnosis of septicemia. Archives of Internal Medicine. 1966;118(4):356.

37. Tang S, Zhang Y, Yin SW, et al. Neutrophil extracellular trap formation is associated with autophagy-related signalling in ANCA-associated vasculitis. Clinical & Experimental Immunology. 2015;180(3):408–418.

38. Elmore SP, Qian T, Grissom SF, Lemasters JJ. The mitochondrial permeability transition initiates autophagy in rat hepatocytes. Faseb Journal Official Publication of the Federation of American Societies for Experimental Biology. 2001;15(12):2286.

39. Welchman RL, Gordon C, Mayer RJ. Ubiquitin and ubiquitin-like proteins as multifunctional signals. Nature Reviews Molecular Cell Biology. 2005;6(8):599–609.

40. Orenstein SJ, Cuervo AM. Chaperone-mediated autophagy: Molecular mechanisms and physiological relevance. Seminars in Cell & Developmental Biology. 2010;21(7):719–726.

41. Zuckerfranklin D, Hirsch JG. ELECTRON MICROSCOPE STUDIES ON THE DEGRANULATION OF RABBIT PERITONEAL LEUKOCYTES DURING PHAGOCYTOSIS. Journal of Experimental Medicine. 1964;120(4):569.

42. Borregaard N, Lollike K, Kjeldsen L, et al. Human neutrophil granules and secretory vesicles. European Journal of Haematology. 1993;51(4):187.

43. László L, Doherty FJ, Watson A, et al. Immunogold localisation of ubiquitin-protein conjugates in primary (azurophilic) granules of polymorphonuclear neutrophils. Febs Letters. 1991;279(2):175–178.

44. Lominadze G, Powell DW, Luerman GC, Link AJ, Ward RA, McLeish KR. Proteomic analysis of human neutrophil granules. Molecular & Cellular Proteomics. 2005;4(10):1503–1521.

45. Berrier AL, Yamada KM. Cell–matrix adhesion. Journal of Cellular Physiology. 2010;213(3):565–573.

46. Borregaard N. Neutrophils, from marrow to microbes. Immunity. 2010;33(5):657–670.

47. Tuloup-Minguez VR, Hamaã A, Greffard A, Nicolas VR, Codogno P, Botti JL. Autophagy modulates cell migration and Î²1 integrin membrane recycling. Cell Cycle. 2013;12(20):3317–3328.

48. Briggs RT, Drath DB, Karnovsky ML, Karnovsky MJ. Localization of NADH oxidase on the surface of human polymorphonuclear leukocytes by a new cytochemical method. Journal of Cell Biology. 1975;67(3):566–586.

49. Rikihisa Y. Glycogen autophagosomes in polymorphonuclear leukocytes induced by rickettsiae. Anatomical Record-advances in Integrative Anatomy & Evolutionary Biology. 1984;208(3):319–327.

50. Quie PG, White JG, Holmes B, Good RA. In vitro bactericidal capacity of human polymorphonuclear leukocytes: diminished activity in chronic granulomatous disease of childhood. Journal of Clinical Investigation. 1967;46(4):668–679.

51. Matsuda N, Sato S, Shiba K, et al. PINK1 stabilized by mitochondrial depolarization recruits Parkin to damaged mitochondria and activates latent Parkin for mitophagy. Journal of Cell Biology. 2010;189(2):211–221.

52. Zucker-Franklin D, Hirsch JG. Electron microscope studies on the degranulation of rabbit peritoneal leukocytes during phagocytosis. The Journal of experimental medicine. 1964;120(4):569.

53. Borregaard N, Lollike K, Kjeldsen L, et al. Human neutrophil granules and secretory vesicles. European journal of haematology. 1993;51(4):187–198.

54. Papadopoulos C, Meyer H. Detection and clearance of damaged lysosomes by the endo-lysosomal damage response and lysophagy. Current Biology. 2017;27(24):R1330–R1341.

55. Bhattacharya A, Wei Q, Shin JN, et al. Autophagy is required for neutrophil-mediated inflammation. Cell reports. 2015;12(11):1731–1739.

56. Haglund K, Dikic I. Ubiquitylation and cell signaling. Embo Journal. 2005;24(19):3353–3359.

57. Ohtake F, Baba A, Takada I, et al. Dioxin receptor is a ligand-dependent E3 ubiquitin ligase. Nature. 2007;446(7135):562–566.

58. Lobert VH, Brech APedersen NM, Wesche J, Oppelt A, Malerod L, Stenmark H. Ubiquitination of alpha 5 beta 1 integrin controls fibroblast migration through lysosomal degradation of fibronectin-integrin complexes. Developmental Cell. 2010;19(1):148.

59. Sturany S, Van JL, Gilchrist A, Vandenheede JR, Adler G, Seufferlein T. Mechanism of activation of protein kinase D2(PKD2) by the CCK(B)/gastrin receptor. Journal of Biological Chemistry. 2002;277(33):29431.

60. Eisenberglerner A, Kimchi A. PKD is a kinase of Vps34 that mediates ROS-induced autophagy downstream of DAPk. Cell Death & Differentiation. 2012;19(5):788–797.

